# A Distinct Phenotype of Polarized Memory B cell holds IgE Memory

**DOI:** 10.1101/2023.01.25.525495

**Authors:** Joshua F.E. Koenig, Niels Peter H Knudsen, Allyssa Phelps, Kelly Bruton, Ilka Hoof, Gitte Lund, Danielle Della Libera, Anders Lund, Lars Harder Christensen, David R. Glass, Tina Walker, Allison Fang, Susan Waserman, Manel Jordana, Peter S Andersen

## Abstract

Allergen-specific IgE antibodies mediate allergic pathology in diseases such as allergic rhinitis and food allergy. Memory B cells (MBCs) contribute to circulating IgE by regenerating IgE-producing plasma cells upon allergen encounter. We report a population of type 2 polarized MBCs defined as CD23^hi^, IL-4Rα^hi^, CD32^low^ at the transcriptional and surface protein levels. These “MBC2s” are enriched in IgG1 and IgG4-expressing cells, while constitutively expressing germline transcripts for IgE. Allergen-specific B cells from patients with allergic rhinitis and food allergy were enriched in MBC2s. MBC2s generated allergen specific-IgE during sublingual immunotherapy, thereby identifying these cells as the primary reservoir of IgE. The identification of MBC2s provides insights into the maintenance of IgE memory, which is detrimental in allergic diseases, but which could be beneficial in protection against venoms and helminths.

**One-Sentence Summary:** Identification of a novel memory B cell subset which holds allergen specific IgE memory.

## Main Text

Allergies to innocuous environmental and food antigens affect >10% of the population and, in some instances, can elicit life-threatening reactions such as anaphylaxis. Aberrant immune responses against allergens are mediated by a type 2-dominant T cell signature that drives IgE production. Some allergies persist for years or even a lifetime as a result of long-lived immunological memory. In recent years, multiplex profiling of allergen specific CD4 T cells has uncovered substantial heterogeneity, facilitating the discovery of Th2 and follicular T helper cell subsets unique to allergic disease (*1–3*). Similar efforts on profiling memory B cells (MBCs) outside of allergy have made apparent that MBCs are not a uniform population, where varied phenotypes provoke specific fates upon reactivation (*4–6*).

A clear consensus has emerged that allergen-specific B cell memory is held by non-IgE MBCs (*7*–*9*). In both mice and humans, the allergen-specific MBC repertoire is dominated by IgG1, while IgE B cells are short-lived and IgE MBCs are exceptionally rare or absent (*8, 10–12*). IgG1-expressing MBCs become IgE-producing plasma cells (PC) during a memory response to allergen exposure, contributing to the persistence of IgE-mediated allergies (*9, 13–16*). IgG1, however, is present in various contexts of human health and disease, including protective immunity against pathogens. IgG1 against allergens is produced by non-allergic individuals, though most will never progress to pathogenic IgE production (*17*). MBC isotype alone is, therefore, not sufficient to explain the capacity of IgG1 MBCs to hold IgE memory. Recent work has suggested that MBC phenotype, rather than isotype, defines the function of MBCs (*5*). The phenotype of the MBCs which hold the memory of IgE responses is poorly characterized.

Here, we generated a single-cell RNA-sequencing (scRNA-seq) atlas of isotype switched MBCs from allergic and non-allergic donors to characterize the MBC repertoire. We identified a novel subset of type 2-polarized MBC, hereafter referred to as “MBC2”, distinguishable by their expression of CD23 (*FCER2*)*, IL4R, IL13RA1*, IgE germline transcripts (εGLT), and the absence of CD32 (*FCGR2B*). MBC2s could also be distinguished based on their surface protein expression phenotype assessed by mass cytometry. We found that human and mouse allergen-specific MBCs adopted an MBC2 phenotype while virus-specific MBCs did not. Furthermore, *in vivo* human memory IgE responses to sublingual allergen immunotherapy were clonally derived from MBC2s. The production of MBC2s was absent in mice lacking the canonical type 2 cytokine IL-4, which demonstrates the close association between MBC2s and allergic immunity. Identification of a type 2 polarized phenotype of MBC has significant implications for the development of allergic disease and immunity and is therefore critical in the design of novel therapeutics in IgE-mediated diseases, including aeroallergies, asthma, and food allergies.

## Results

### scRNA-seq atlas of switched memory B cells reveals a distinct population of MBCs expressing markers of type 2 polarization

To interrogate the phenotype of MBCs in allergic subjects, we performed scRNA-seq on isotype switched MBCs purified by negative selection from the peripheral blood mononuclear cells (PBMCs) from 6 birch-allergic subjects at outside of the birch season, 4 house dust mite allergic subjects, and 5 non-allergic controls (Fig. 1A). Following pre-processing (Methods), the final data set included 90,418 human MBCs. Unsupervised clustering revealed 21 clusters of MBCs, many of which formed distinct “islands” when projected in two dimensions. (Fig. S1A). All MBC clusters were represented in PBMCs from allergic and non-allergic subjects with similar frequency indicating that allergic status is not defined by a single population of circulating MBC (Fig. S1B, C).

**Fig. 1:**
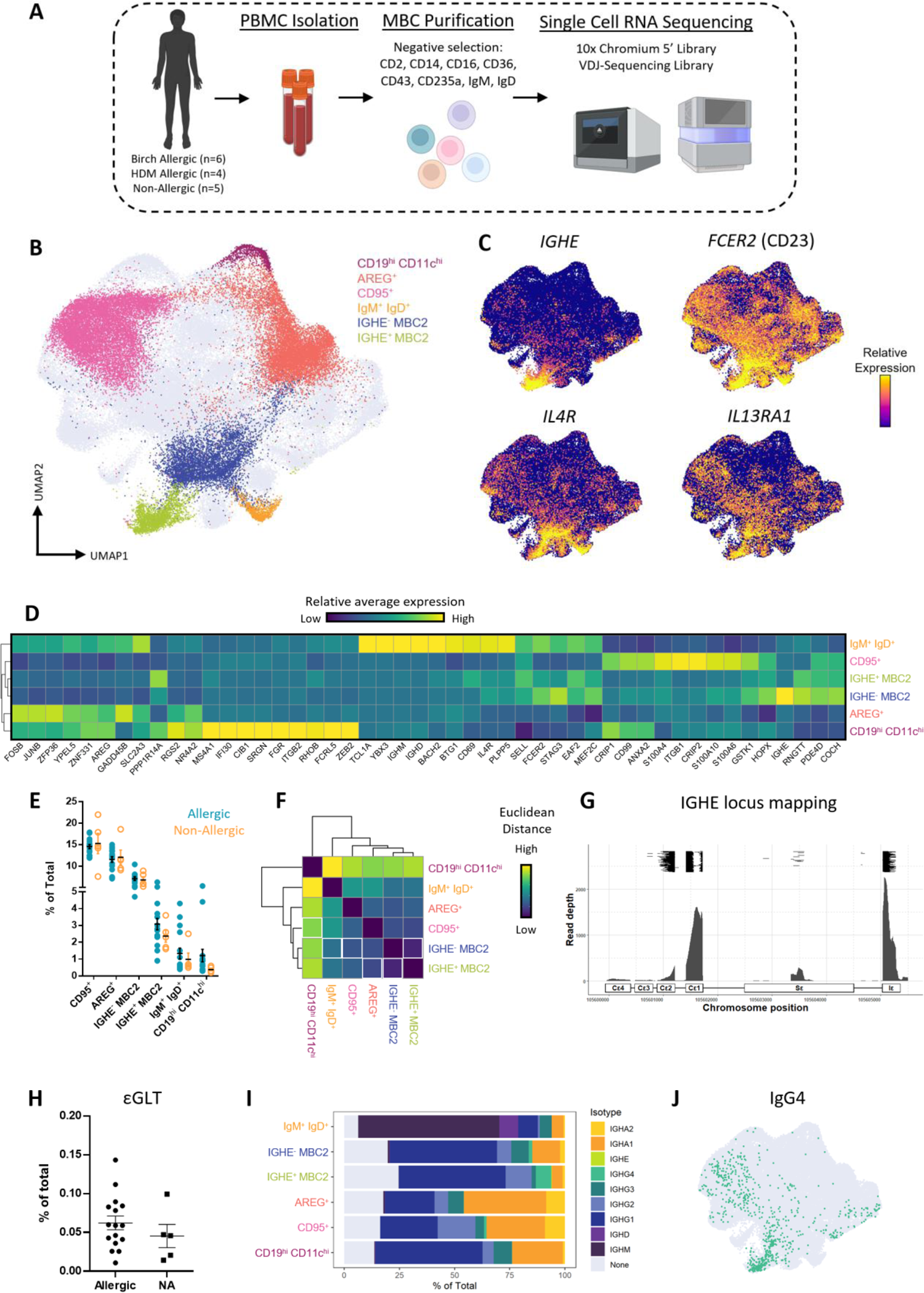
scRNA-seq atlas of switched memory B cells revealed a distinct population of MBCs expressing markers of type 2 polarization. (**A**) Schematic of experimentation. MBCs were negatively selected from PBMCs from birch-, house dust mite (HDM)-, and non-allergic individuals and single cell sequenced. (**B**) UMAP plot of MBC clusters. Four clusters were selected for comparison to two type 2 polarized MBC (MBC2) clusters. Analysis of all clusters is presented in Figures S1 and S2. (**C**) UMAP plot colorized by markers associated with type 2 immunity. (**D**) Heatmap of scaled average expression of the top 10 differentially expressed genes for each cluster. (**E**) Frequency of each cluster among total MBCs per participant. (**F**) Euclidean distances between clusters based on mean expression. Lowest Euclidean distance per cluster is denoted with a white outline. (**G**) Mapping of IGHE reads onto the IgH locus on chromosome 12. Black lines at the top represent paired-end reads. (**H**) Frequency of all MBCs which express εGLT in allergic patients and non-allergic controls. (**I**) Isotype identified in antibody encoding reads from the VDJ sequencing library. (**J**) UMAP colorized by IgG4-expression based on VDJ sequencing library.

We identified two novel clusters of MBCs that highly express markers associated with type 2 immunity, which we have denoted “MBC2”. (Fig. 1B, C). The unifying features of these subsets were high expression of the low-affinity IgE receptor CD23 (*FCER2*), and receptors for the type 2-associated cytokines IL-4 (*IL4R*) and IL-13 (*IL13RA1*). The two MBC2 clusters differed primarily by the expression of reads which map to the *IGHE* locus in one cluster (IGHE^+^ MBC2) and the absence of IGHE and greater *CD69* expression in the other (IGHE-MBC2). When *IGHE* was not considered in the clustering analysis, these two populations co-clustered, suggesting that they are transcriptionally related (Fig. S1D).

To contextualize and compare the MBC2 cluster with other MBCs, we selected four additional clusters based on existing literature and their distribution in UMAP space (Fig. S1E; A discussion of all 21 clusters is present in the *Supplemental Text* and analyses including all clusters are found in Fig. S2). These four selected clusters include: [1] a cluster of *IGHM*^+^ *IGHD*^+^ *CD27*^-^ cells, a phenotype consistent with naïve B cells which bypassed the negative selection kit; [2] a cluster of cells characterized by high expression of *CD19*, *ITGAX* (CD11c), and *FCRL5* which are referred to as atypical B cells or age-associated B cells, but are denoted here as “CD19^hi^ CD11c^hi^” (*5, 18*); [3] a subset identified in a recent surface marker screen of MBCs by Glass *et al.* which are CD95^+^ *FAS^+^* (CD95), *ITGB1^+^* (CD29), *NT5E^+^* (CD73) cells denoted as “CD95^+^” (*5*), and [4] a cluster of MBCs expressing *AREG, CDKN1A,* and *NFKB1*, a phenotype similar to that reported by bulk RNA-sequencing of citrullinated peptide-specific B cells in rheumatoid arthritis patients (denoted “AREG^+^”) (*19*). These selected clusters were present at similar frequencies in both allergic and non-allergic subjects (Fig. 1E).

Each of the six MBC clusters had defining transcriptional characteristics (Fig. 1D). The IGHE^+^ and IGHE^-^ MBC2 clusters shared a largely overlapping transcriptional profile. This profile was unique compared to the other clusters (Fig. 1D, Fig. S2B). Indeed, we calculated the Euclidean distance between clusters considering all transcribed genes and found that the IGHE^+^ and IGHE^-^ MBC2 clusters were highly similar and were more distantly related to other MBC clusters (Fig. 1F, Fig. S2C). The two MBC2 clusters shared high *IL4R* and *FCER2* expression with IgM^+^ IgD^+^ cells, but these genes were downregulated in the other MBC subsets. These IgM^+^ IgD^+^ cells expressed genes associated with naïve B cells (*TCL1A, BACH2, IGHM, IGHD)*. CD95^+^ MBCs differentially expressed *CRIP1, CRIP2, and ITGB1* (CD29), but shared expression of the transcription factor *HOPX* with MBC2s. AREG^+^ MBCs differentially expressed markers associated with repression of activation (*e.g., NR4A2*, *GADD45B*, *ZFP36*). CD19^hi^ CDl1c^hi^ MBCs had the greatest Euclidean distance to any other MBC subset, and shared expression of very few differentially expressed genes with the MBC2 clusters (Fig. S2E). Interestingly, while the MBC2 populations expressed markers associated with type 2 immunity, CD19^hi^ CD11c^hi^ MBCs expressed markers associated with type 1 immunity, including *TBX21* (Tbet)(*20*), *ZEB2* (*21*), and evidence of IFN-γ stimulation (*e.g.*, *IFI30*).

The IGHE-expressing MBCs were present at a much greater frequency in our scRNA-seq dataset than previous careful reports of the frequency of IgE MBCs(*8*). We, therefore, mapped the raw sequencing reads to the IGHE locus and found that these reads mapped to the Iε and ε switch region in addition to the ε constant region, indicating that these are εGLTs rather than productively rearranged IgE transcripts (Fig. 1G). Indeed, the majority of IGHE^+^ MBC2s cells co-expressed another immunoglobulin isotype, primarily IGHG1 (Fig. 1I). The frequency of εGLT-expressing cells was consistent between individuals regardless of allergic status (Fig. 1H). IGHE^+^ MBC2s are therefore a physiological subset of MBCs expressing εGLT and are not a correlate of allergic disease.

Analysis of Ig constant regions from the amplified VDJ library revealed that IGHE^+^ and IGHE^-^ MBC2 clusters were enriched in IgG1 and IgG4 isotype expression and had the lowest expression of IgA isotypes among switched MBCs (Fig. 1I, Fig. S2D). IgG1 and IgG4 are both induced by IL-4, further associating the MBC2 clusters with type 2 immunity (*22–24*). In contrast, CD95^+^ and AREG^+^ MBCs had a high frequency of IgA1 and IgA2 expression, and greater representation of IgG2 and IgG3 (Fig. 1I, Fig. S2D). CD19^hi^ CD11c^hi^ MBCs were also enriched in IgG1^+^ cells and had the highest representation of IgG3^+^ cells (Fig. 1I, Fig. S2D). Only 8 of the 90,418 cells expressed a productively rearranged IgE antibody, all of which clustered with the IGHE^+^ MBC2s (Fig. 1I, Fig. S2D). Collectively, the transcriptional and isotype expression profiles of MBC2s are consistent with type 2 polarization.

### MBC2s have a distinct surface phenotype

We next sought to evaluate whether MBC2 cells could be distinguished based on their surface protein expression using a mass cytometry dataset from Glass *et al.* designed to characterize the diversity of human B cells in various tissues (*5*). Since εGLT expression cannot be evaluated at the protein level, we focused on CD23 as a primary marker of MBC2s. CD23 expression is thought to be downregulated as naïve B cells are activated and transition into a memory fate (*6*). Using the annotations from Glass *et al.* we depleted naïve B cells *in silico* and observed that surface CD23 was retained by ∼5% of non-naïve IgD^-^ B cells. This frequency was approximately consistent with the MBC2 phenotypes observed by scRNA-seq (Fig. 1E, Fig. 2A). CD23^+^ MBCs had variable expression of the MBC markers CD27 and CD45RB (Fig. 2A). High-dimensional analysis revealed MBC2s as a distinct population with a unique surface expression pattern compared to the other MBC phenotypes described by Glass *et al.* (Fig. 2B–E, Fig. S3A). MBC2 frequency was highest in circulation, but they could also be detected in the tonsil, bone marrow, and lymph nodes, where they existed with minor tissue-specific variations in expression (Fig. S3B–C).

**Fig. 2:**
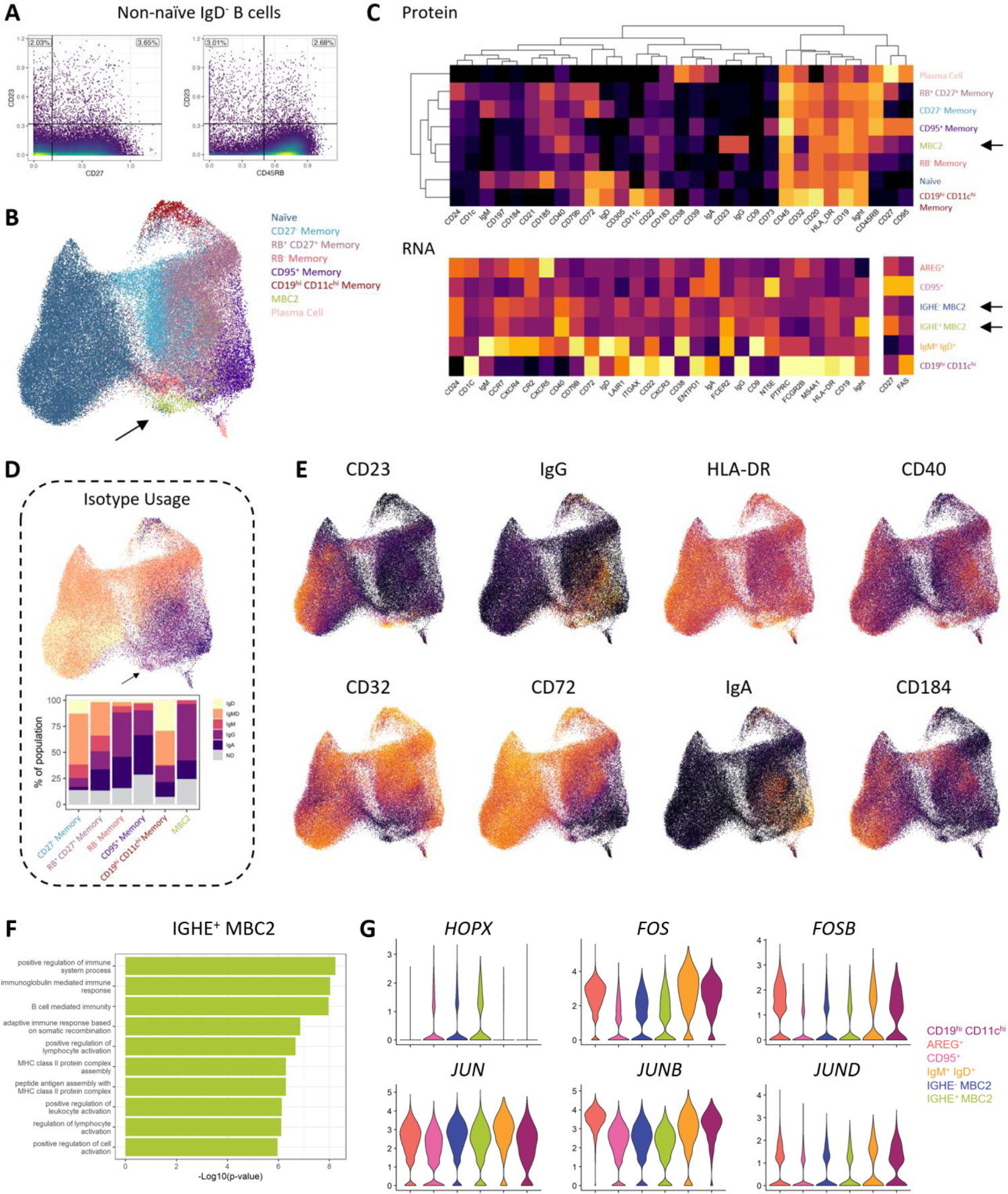
MBC2s have a distinct surface phenotype. (**A**–**E**) Re-analysis of high-dimensional human MBCs mass cytometry dataset from *Glass et al*. (A) CD23 expression on IgD^-^ non-naïve B cells. (B) UMAP plot of B-lineage cells. (C) Top: Heatmap of median surface protein expression by subset from mass cytometry data. Bottom: Heatmap of scaled mean RNA-expression of markers included in the mass cytometry dataset per cluster. The same markers are aligned in columns, labels in the top heatmap are in protein annotation format whereas labels in the bottom heatmap are in gene annotation format. (D) Isotype usage by subset; ND = “not determined”, IgMD = co-expression of IgM and IgD. (E) UMAP colorized by marker expression. (**F**–**G**) Analysis of MBC2 gene expression from scRNA-seq atlas. (**F**) Top 10 Gene Ontology terms for upregulated differentially expressed genes in the IGHE^+^ MBC2 population. (**G**) Normalized and scaled RNA expression of HOPX and genes of the AP-1 transcriptional program by cluster.

To further understand the MBC2 phenotype, we compared gene expression from our scRNA-seq dataset with the markers included in the mass cytometry panel, with the exception of CD45RB as the specific exons of CD45 are difficult to evaluate from scRNA-seq data (Fig. 2C). The scRNA-seq and mass cytometry dataset showed remarkable similarity across MBC subsets. At both the RNA and protein levels, MBC2s had elevated levels of CD40, CD23 (*FCER2*), and HLA-DR compared to other subsets. MBC2s also had a unique downregulation of the inhibitory IgG Fc receptor CD32 (*FCGR2B*). IGHE^-^ and IGHE^+^ MBC2s differed in expression of some markers; a granularity that was not discernable within the mass cytometry dataset because of the inability to distinguish the MBC2 clusters by εGLT expression. For example, MBC2s had detectable surface protein expression of CD22, CD72, and CD183 (*CXCR3*), which was primarily transcribed by IGHE^-^ MBC2s. The enrichment in IgG^+^ cells and relative paucity of IgA^+^ cells among MBC2s observed by scRNA-seq was consistent at the level of protein expression (Fig. 2D).

The CD95^+^ and CD19^hi^ CD11c^hi^ clusters identified by scRNA-seq had closely matching gene expression and surface phenotypes to their respective mass cytometry clusters (Fig. 2C–E). Namely, CD95^+^ cells had high expression of CD95 (*FAS*), CD27, CD73 (*NT5E*), CD39 (*ENTPD1*), and low expression of CD23 (*FCER2)*, CD72, CD305 (*LAIR1*), HLA-DR, relative to other subsets. CD19^hi^ CD11c^hi^ cells had high expression of CD19, CD11c (*ITGAX*), CD20 (MS4A1), CD22, CD183 (*CXCR3*), and low expression of CD21 (CR2), CD24, CD40, CD23 (*FCER2)*, CD73 (*NT5E),* CD197 (*CCR7*) compared to other subsets. These data provide additional validation of the MBC nomenclature established by Glass *et al.* Gene ontology analysis of the top differentially expressed genes for IGHE^+^ MBC2 cells revealed an enrichment in pathways associated with adaptive immune activation and antigen presentation via MHC-II (Fig. 2F). Enrichment in these pathways was driven in part by the upregulation of the costimulatory molecule *CD40*, and of various *HLA-DR/DQ* genes (Table S1). The transcription factor *HOPX* was identified as a top differentially expressed gene shared by both MBC2 clusters and CD95^+^ MBCs (Fig. 2G). *HOPX*-expressing MBC clusters had lower expression of JUN and FOS genes (Fig. 2G), in line with a known role for *HOPX* in inhibiting the AP-1 transcriptional program (*25, 26*). Inhibition of the AP-1 transcriptional program is thought to be involved in maintaining the memory state and inhibiting PC differentiation (*27, 28*). Taken together, MBC2s present as a type 2 polarized subset of B cells transcriptionally restricted in a memory state, but which appear poised for activation based on their downregulation of inhibitory receptors, and upregulated antigen-presentation and co-stimulation machinery.

### Human allergen-specific MBCs are primarily MBC2s

Given the type 2-polarized gene expression of the MBC2 phenotype, we hypothesized that allergen-specific MBCs would frequently adopt this phenotype. As allergen-specific B cells are exceedingly rare in whole PBMCs, we used Bet v 1 tetramers to enrich birch-specific cells from patients with allergic rhinitis prior to single sorting MBCs for SMARTSeq2 scRNA-seq library preparation (Fig. 3A, B). In total, 52 tetramer-bound cells were sorted, most of which were found to be allergen specific when expressed as recombinant antibodies (22/30, 67%. Fig. 3C). We supplemented this dataset with previously published transcriptomes from grass-specific MBCs from allergic patients (*16*). For comparison, we sorted MBCs which bound a tetramer of the receptor binding domain (RBD) of SARS-CoV-2, which arise in the context of viral infection and/or vaccination and are therefore primarily driven by type 1 immunity. This approach allowed for a transcriptomic comparison of allergen- and virus-specific MBCs and, further, allowed us to compare the transcriptional signature with cells in our scRNA-seq MBC atlas.

**Fig. 3:**
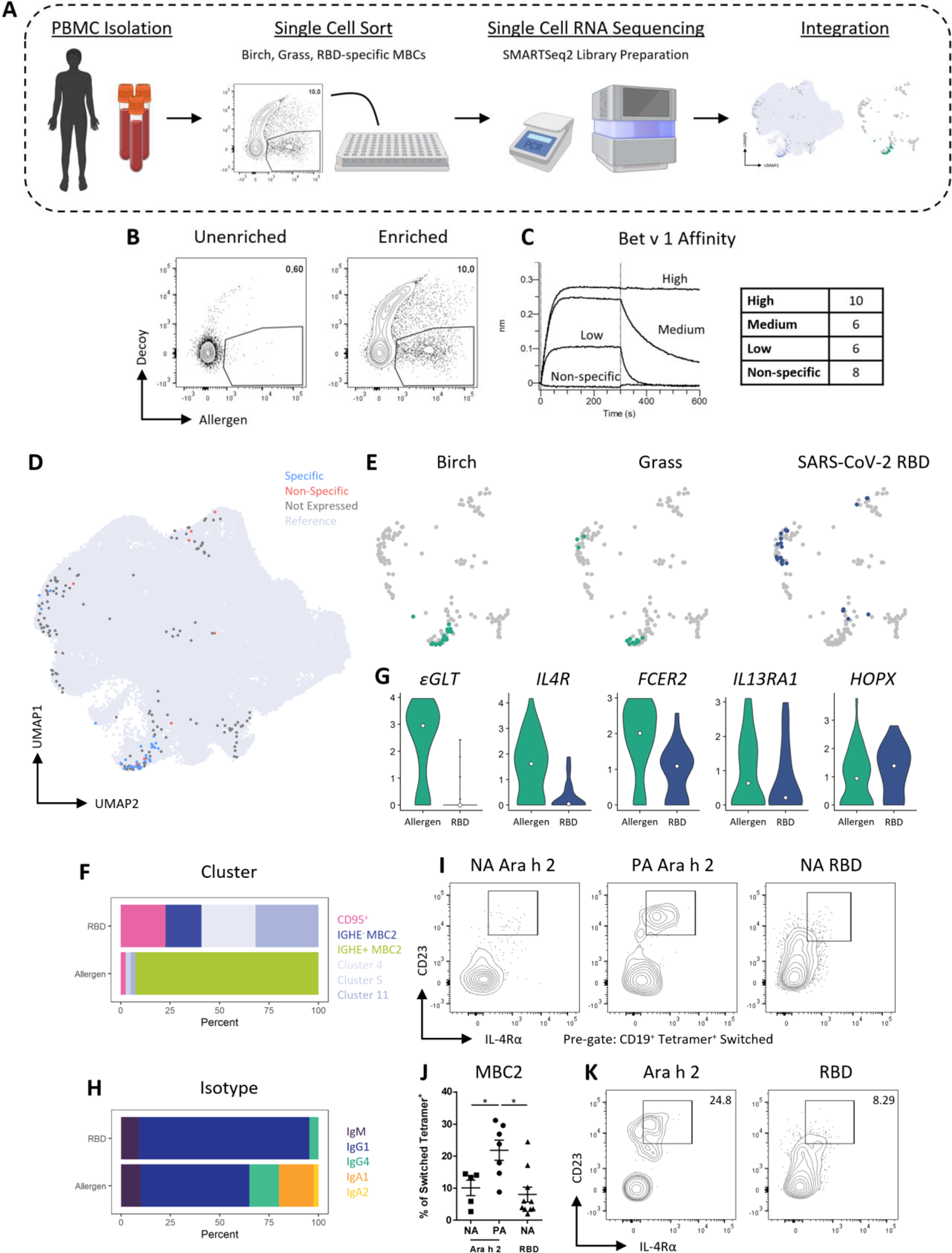
Allergen-specific MBCs are primarily MBC2s. (**A**) Schematic of experimentation. scRNA-seq libraries from single-sorted allergen-specific MBCs were mapped onto the MBC atlas UMAP. (**B**) Sample flow cytometry plot of allergen tetramer-enriched MBCs. (**C**) Allergen-specificity of recombinantly expressed antibodies from Bet v 1-specific MBC transcriptomes evaluated by biolayer interferometry. The number in the table refers to the number of expressed antibodies in each category. (**D**) UMAP plots of integrated transcriptomes from allergen tetramer-binding MBCs colorized by antigen specificity of recombinantly expressed antibodies as measured by ELISA. (**E**) UMAP plots separated by specificity. (**F**) Cluster assignment of single-sorted transcriptomes separated based on allergen- or viral-specificity. (**G**) Normalized and scaled expression of MBC2-associated genes. White dot represents median expression. (**H**) Isotype identified by RNA expression. (**I**–**J**) The phenotype of MBCs specific for the major peanut allergen Ara h 2, or the receptor binding domain (RBD) of native SARS-CoV-2 were evaluated in non-allergic participants (NA) or highly sensitized (peanut-IgE >70 kU/ml) or peanut allergic participants (PA) by flow cytometry. (I) Concatenated flow plots and (J) cell number of tetramer-binding MBC2s. (**K**) Flow cytometry plot of Ara h 2- or RBD-specific MBC2s from pooled PBMCs from 3 PA patients. * p < 0.05 via one-way ANOVA.

We integrated the sorted single cell transcriptomes with transcriptomes from our MBC atlas (Fig. 1). When plotted onto the MBC atlas UMAP, the majority of allergen-specific transcriptomes clustered with MBC2 cells (Fig. 3D, E). In contrast, RBD-specific B cells primarily clustered with MBCs related to CD95^+^ MBCs, revealing an association between the context of immune activation and the polarization of MBCs (Fig. 3 D–F). Indeed, allergen-specific MBCs expressed defining markers of the MBC2 phenotype, while most RBD-specific cells did not (Fig. 3G). Interestingly, both allergen-specific and RBD-specific B cells primarily expressed IgG1, further demonstrating that phenotype rather than isotype is required to distinguish allergic and non-allergic MBC subsets (Fig. 3H). The phenotype of allergen-specific MBCs was confirmed at the protein level and in another allergic context by surface staining for CD23 and IL-4Rɑ on Ara h 2 allergen tetramer-specific B cells from highly sensitized peanut allergic patients (serum peanut-specific IgE > 70 kU/mL) and non-allergic controls. Peanut allergic patients had a two-fold greater population of CD23^hi^ IL-4Rα^hi^ switched B cells which bound to the peanut allergen Ara h 2 compared to non-allergic individuals (Fig. 3I, J). We further compared allergen-specific B cells and virus-specific B cells in pooled PBMCs from three SARS-CoV-2-immunized peanut allergic patients, revealing a clear difference in the frequency of CD23^hi^ IL-4Rα^hi^ B cells between the two repertoires (Fig. 3K). In summary, the presence of allergen specific MBC2s is characteristic of allergic sensitization rather than a tolerant response to allergens and is associated with type 2 polarization rather than type 1 polarization.

### Allergen-specific MBC2s are induced in mouse models of allergy

To facilitate mechanistic experimentation on MBC2 ontogeny, we examined the phenotype of mouse MBCs to determine whether a homologue to the human MBC2 exists. We sensitized separate groups of mice with ovalbumin (OVA) using multiple type 2-immune driven models and routes of administration and evaluated the surface expression of the MBC2 markers CD23 and IL-4Rɑ on OVA-specific MBCs (Fig. 4A). These models included oral sensitization in the presence of the type 2 adjuvant cholera toxin (CT), a model of skin sensitization via tape stripping, and intraperitoneal immunization with CT. As a comparator, we immunized an additional group of mice with OVA using CpG as an adjuvant, a known driver of type 1 immunity. MBC phenotypes were evaluated in the OVA tetramer-binding population following enrichment, the same methodology we used to study human MBCs (Fig. 3A, I). Across type 2 sensitization models, we observed a population of OVA-specific CD23^hi^ MBCs. These cells also had higher expression of IL-4Rα, though all mouse MBCs expressed the receptor (Fig. 4B, D). In contrast, only a small number of non-allergen-specific switched B cells from the same mice were CD23^hi^ IL-4Rα^hi^. (Fig. 4C, D). Further, CD23^hi^ IL-4Rα^hi^ OVA-specific MBCs were nearly absent in mice immunized in type 1 conditions (Fig. 4B, D). We next contextualized the surface phenotype of CD23^+^ IL-4Rα^hi^ cells with previously published phenotypes of MBCs. CD23^hi^ IL-4Rα^hi^ cells expressed CD80, CD73, and PD-L2, a phenotype associated with MBC maturity and potent IgE production during memory responses (Fig. S4)(*4, 9*).

**Fig. 4:**
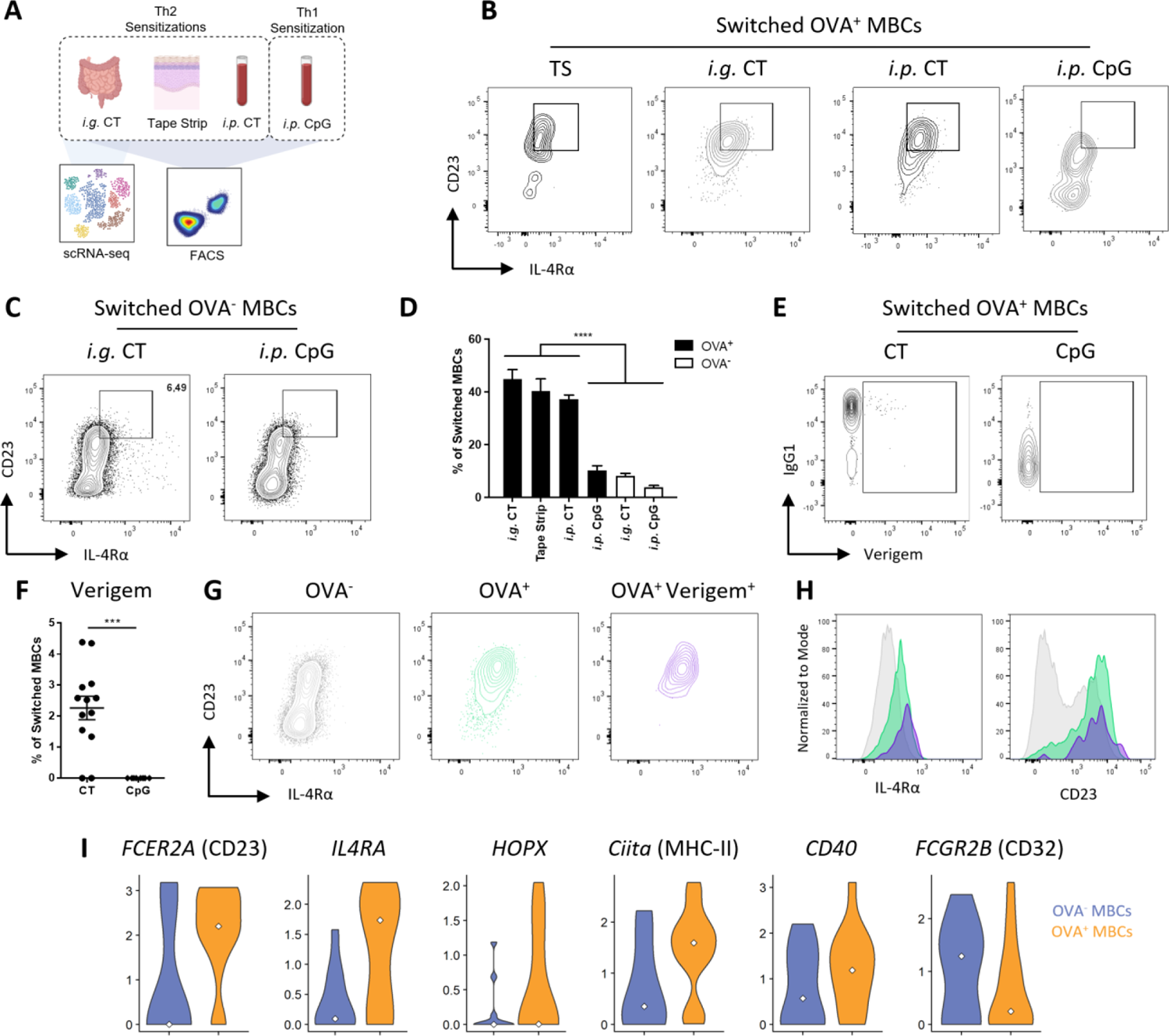
Identification of mouse MBC2s. (**A**) Schematic of contexts in which MBC phenotypes were evaluated using scRNA-seq or flow cytometry. (**B**–**C**) Representative flow cytometry plots of surface CD23 and IL-4Rα expression on (B) OVA^+^ or (C) non-allergen-specific switched MBCs (IgM^-^ IgD^-^ CD38^+^ GL7^-^). (**D**) Summary plots of the frequency of MBC2s among switched MBCs. (**E**) Concatenated plots of IgG1 and Verigem expression on OVA-specific MBCs from *i.p.* OVA+CT-immunized mice. (**F**) Summary plot of Verigem^+^ MBCs from switched MBCs. (**G**) Concatenated flow cytometry plots of CD23 and IL-4Rα on OVA^-^, OVA^+^, and OVA^+^ Verigem^+^ switched MBCs. (**H**) Histograms of IL-4Rα and CD23 on subpopulations of switched MBCs of populations from (G). (**I**) RNA expression of MBC markers from single-sorted OVA-specific MBCs from OVA+CT immunized mice. n=5-13, minimum of 2 independent experiments per treatment condition. * p < 0.05 ** p < 0.01 *** p < 0.001 **** p < 0.0001 via Mann-Whitney test when comparing two groups or one-way ANOVA when comparing 6 groups.

We next sought to detect εGLT, the most distinguishing feature of human MBC2s, *in vivo* using Verigem mice which have a YFP reporter inserted downstream of the IgE constant region. YFP should be produced when the full length εGLT is transcribed. ∼2% of OVA-specific MBCs expressed Verigem, the majority of which co-expressed IgG1 (Fig. 4E, F). In contrast, OVA-specific MBCs from Verigem mice immunized with CpG did not have detectable Verigem expression (Fig. 4E, F). These Verigem^+^ OVA-specific MBCs expressed high levels of CD23 and IL-4Rα, consistent with MBC2 polarization (Fig. 4G, H). We further validated the MBC2 phenotype by single-sorting OVA-specific and non-OVA-specific switched MBCs and single cell sequencing them using a SMARTSeq2 library preparation. Mouse OVA-specific MBCs had a similar gene expression profile as human MBC2s; greater transcript expression of CD23, IL-4Rα, HOPX, MHC-II, CD40 and lower expression of CD32 than non-OVA-specific MBCs (Fig. 4I). While mouse MBCs do not express εGLT at the same frequency as human MBCs, these cells share many transcriptional and surface protein expression features. We have, therefore, identified and characterized the conserved phenotype of MBC2s within the human and murine MBC repertoire.

### MBC2 generation does not require germinal centers

A recent study reported the existence of CD23^+^ IL-4Rα^+^ germinal center (GC) B cells which were concluded to be a precursor to MBCs (*29*). Given the similarity of this phenotype to MBC2s, we sought to determine whether MBC2 polarization required passage through the GC. We performed type 2-driven immunizations of B cell-conditional Bcl6 knockout mice (Mb1^Cre/+^ Bcl6^fl/fl^, referred to as GC KO). As previously described, these mice were deficient in GC B cells (Fig. 5A). GC KO mice generated far fewer switched OVA-specific MBCs compared to control (Mb1^+/+^ Bcl6^fl/fl^) mice (Fig. 5B, C). OVA^+^ MBCs from GC KO mice had increased expression of CD23 and IL-4Rα compared to non-antigen-specific MBCs from the same mice (Fig. 5D). However, the frequency and number of OVA-specific MBC2s was significantly reduced compared to wild type (WT) mice (Fig. 5E). MBC2 polarization can, therefore, occur independent of the GC, while MBC2 number is greater with intact GCs.

**Fig. 5:**
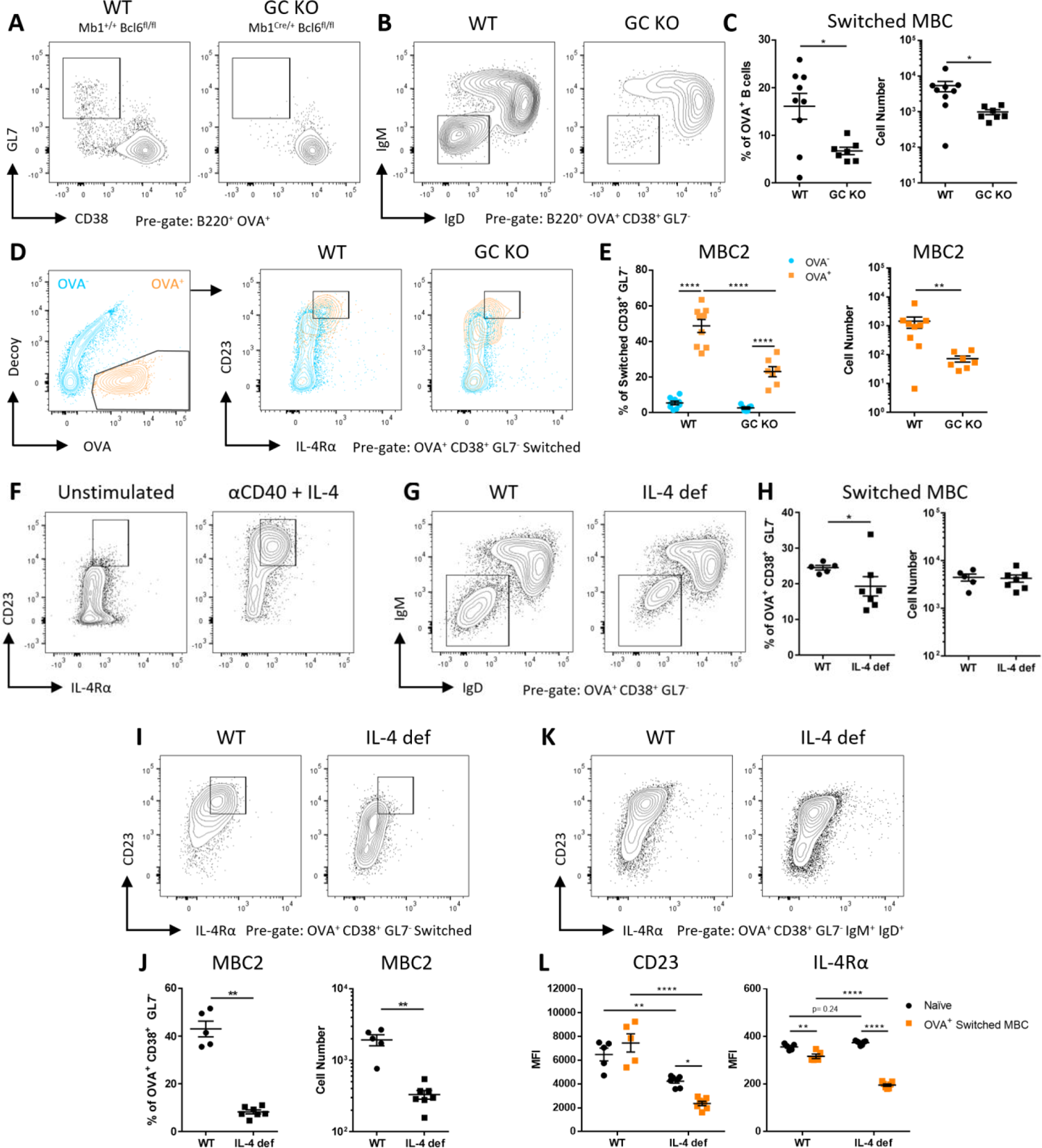
IL-4, but not the GC, is required for MBC2 differentiation. (**A**–**F**) Mb1^+/+^ Bcl6^fl/fl^ (WT) and Mb1^Cre/+^ Bcl6^fl/fl^ (GC KO) were immunized intraperitoneally with OVA and CT and pooled splenocytes and mesenteric lymph node cells were analyzed by flow cytometry at 1-month post-immunization. (A-B) Representative flow plots of (A) OVA-specific GC B cell and (B) MBC isotype. (C) Frequency and number of IgM^-^ IgD^-^ CD38^+^ GL7^-^ OVA-specific MBCs. (D) Representative flow plots of OVA-specific MBC2s. (E) Frequency and number of OVA-specific MBC2s. (F) Splenocytes from naïve mice were stimulated with anti-CD40 and IL-4 for 5 days. CD23 and IL-4Rα expression was evaluated on CD19^+^ CD38^+^ cells. (**G**–**K**) IL-4 deficient (IL-4 def) and WT mice were sensitized i.p. with OVA and CT. (G–H) Concatenated flow plots, frequency and cell number of switched OVA-specific MBCs from pooled spleen and mesenteric lymph nodes. (I–J) Concatenated flow plots, frequency, and cell number of OVA-specific switched MBC2s. (K) Concatenated plots of expression of CD23 and IL-4Rα on naïve B cells. * p < 0.05 ** p < 0.01 *** p < 0.001 **** p < 0.0001 via Mann-Whitney test when comparing two groups or two-way ANOVA when comparing 4 groups. n = 5-8 per group, two independent experiments.

### IL-4 is required for MBC2 differentiation, while secreted IgE is redundant

Since MBC2s are associated with allergen-specific B cells, IgG1, IgG4, and εGLT, we sought to determine whether their production relied on the canonical type 2 cytokine IL-4. To determine whether IL-4 drives differentiation to an MBC2-like phenotype, we stimulated splenocytes from non-allergic mice *in vitro* with anti-CD40 and IL-4 and observed an upregulation of both CD23 and IL-4Rα on switched CD38^+^ memory-like B cells compared to unstimulated cultures (Fig. 5F). To determine if IL-4 is required for MBC2 differentiation *in vivo,* we sensitized mice deficient in IL-4 due to a homozygous replacement of the endogenous IL-4 locus with human CD2 (*30*). IL-4-deficient mice generated a similar number of switched OVA-specific MBCs as WT mice, although there was a lower frequency of switched cells among OVA^+^ CD38^+^ cells (Fig. 5G, H). Among these switched MBCs, IL-4 deficient mice had a near complete absence of CD23^hi^ IL-4Rα^hi^ MBC2s (Fig. 5I, J). The absence of MBC2s was not due to a homeostatic downregulation of IL-4Rα as naïve B cells from IL-4 deficient mice had a similar expression of both IL-4Rα and CD23 compared to WT mice (Fig. 5K, L). IL-4 is therefore a critical requirement for MBC2 differentiation.

IgE receptors are known to be upregulated by the presence of circulating IgE and, therefore, the upregulation of CD23 observed on MBC2s could be related to increases in circulating IgE upon sensitization (*31–33*). We therefore sensitized IgE deficient mice and WT controls and quantified the frequency of MBC2s and their expression of CD23. IgE deficient mice had a similar frequency and number of MBC2s compared to WT controls (Fig. S5A–B). MBC2s from IgE-deficient mice expressed slightly less surface CD23, consistent with receptor downregulation in the absence of IgE (Fig. S5C). Thus, neither the differentiation nor the phenotype of MBC2s rely on the presence of circulating IgE.

### Human MBC2s generate memory IgE responses in vivo

We next sought to determine the functional importance of MBC2s in holding the memory of IgE responses. Circulating IgE is produced by IgE PCs, which are generated from MBCs following allergen exposure (*9, 16, 34*). Addressing the connection between human MBCs and IgE PCs is challenging because both populations are exceedingly rare and, typically, there is a lack of temporal control of allergen exposure in humans. We, therefore, sought to address whether MBC2s are a reservoir of memory IgE responses in a human experimental system using a cohort of birch allergic patients undergoing sublingual immunotherapy (SLIT). The first month of SLIT treatment is known to robustly induce allergen specific IgE from memory which then declines over the course of therapy, as tolerance sets in (*35*). We utilized the controlled induction of high levels of IgE mRNA in blood by sublingual high-dose allergen exposure to determine the clonal connections between MBCs and IgE from PCs at one month of SLIT (*16*).

We amplified bulk VDJ sequences from the PC-containing fraction of enriched PBMC samples collected at one month of SLIT (Fig. 6A). Our MBC atlas contained cells from birch allergic participants at baseline and 1 month of SLIT, allowing us to form clonal lineage connections between the amplified IgE VDJs and the MBCs in our atlas. We were able to identify 52 MBCs that were clonally related to IgE VDJ sequences at 1 month of SLIT (Fig. 6B). 71% of IgE-related MBCs fell within one of the two MBC2 clusters (Fig. 6C); most of which clustered with IGHE^+^ MBC2s (35/52 IGHE^+^, 2/52 IGHE^-^). IgE-related cells were identified both from the baseline and one-month SLIT timepoints, which is consistent with simultaneous IgE PC differentiation and expansion of allergen-specific MBC2s during a memory response (Fig. 6D). Approximately 75% of the clonally related cells expressed IgG (Fig. 6E). Among IgG isotypes, IgG1 and IgG2 were highly represented consistent with previous literature on the isotype switching origins of IgE (*14*). 78% of the IgE-related MBCs were specific for birch extract or Bet v 1, even though the VDJ sequences were not selected based on antigen-specificity (Fig. 6F). These data demonstrate how unlikely a connection between MBCs and circulating IgE is at baseline. It is only when memory is driven by allergen exposure, in this case due to SLIT, that connections between the IgE and MBC2 repertoires can be identified.

**Fig. 6:**
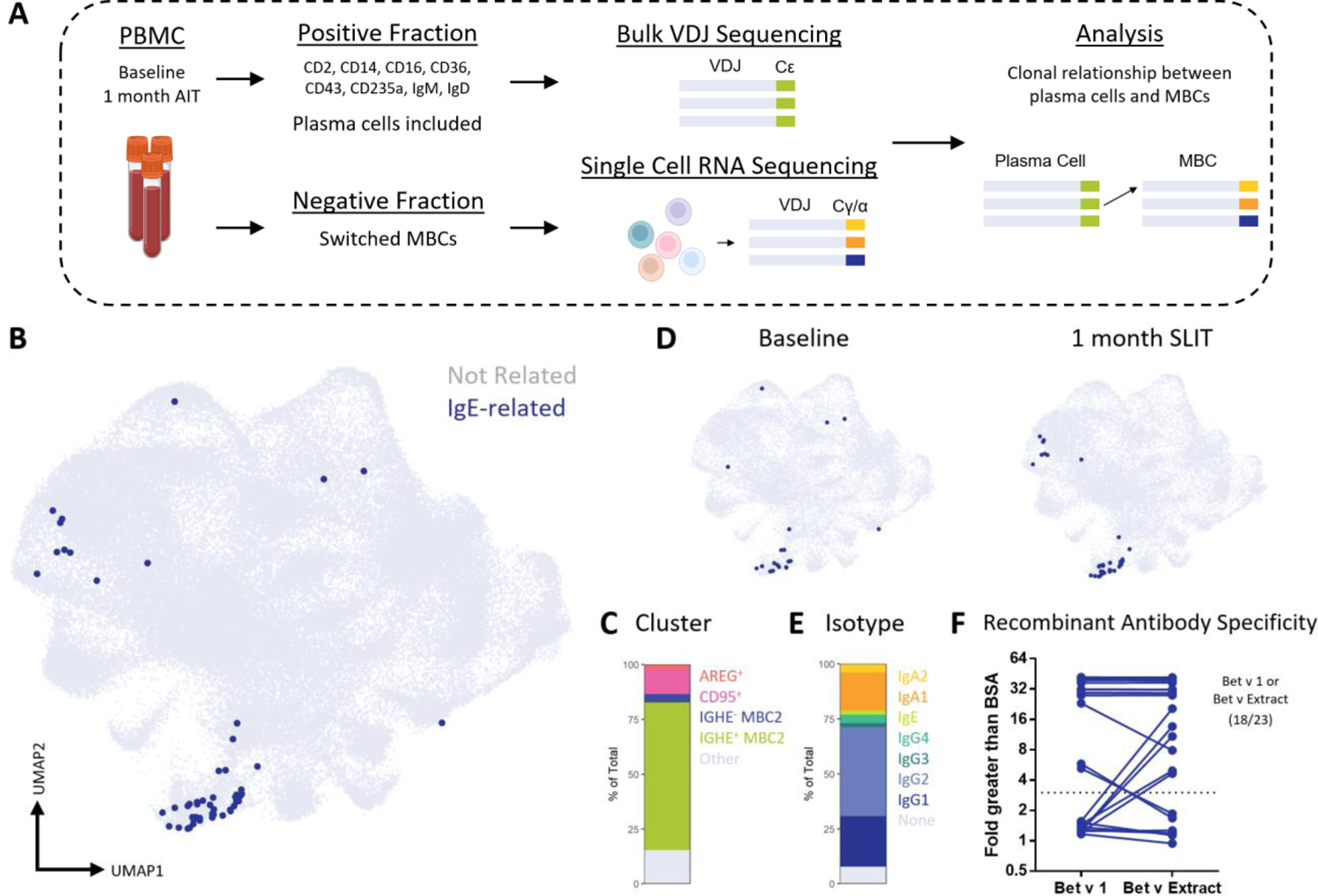
Human MBC2s generate memory IgE responses *in vivo.* VDJ sequences from IgE antibodies were sequences from the plasma cell-containing fraction of MBC purification from 1 month of SLIT and clonally related MBCs were identified from the MBC atlas (**A**) Schematic of the experiment. (**B**) UMAP plot with IgE clonal relatives highlighted. (**C**) UMAP plot separated by SLIT timepoint. (**D**) Antigen-specificity of recombinantly expressed antibodies from IgE-related MBCs evaluated by ELISA. (**E**) Isotype of IgE-related MBCs identified in antibody encoding reads from the VDJ-sequencing library. (**F**) Cluster of IgE-related MBCs.

## Discussion

Over the past decade, it has become increasingly apparent that IgE immunity is not maintained by IgE^+^ MBCs, but instead by upstream isotypes (*7–9, 14, 16, 34*). Despite these advances, the functional properties of MBCs that permit IgE switching remain unresolved, thus curbing the development of novel therapeutics and biomarkers of allergic disease. In an effort to annotate the cellular characteristics that define IgE memory, we produced a comprehensive single-cell transcriptional atlas from over 90,000 human MBCs from patients with IgE-mediated allergies and healthy controls. Our work revealed a distinct type 2-polarized MBC population that distinguishes allergen-specific memory.

MBC2 are defined by high expression of CD23 and IL-4Rα, expression of HOPX and IL-13RA1, and a lack of CD32 expression. Both mouse and human MBC2s express εGLT, though this expression is more frequent in human MBC2s. Expression of CD23, IL-4Rα, and the IGHE locus are under the control of IL-4 (*36–39*); thus, the continued high expression of these markers indicates that resting MBC2s retain IL-4-driven polarization. We have confirmed previous *in vitro* observations that *IL*-4 stimulation drives CD23 and IL-4Rα expression in B cells (*37, 39*). We expanded upon these observations by demonstrating that MBC2s are absent in IL-4-deficient mice, indicating the critical role of IL-4 in MBC2 generation. CD23 expression correlates with allergen specific IgE in allergic patients (*32*). CD23 is reported to enhance antigen-presentation, and co-aggregates with CD21 on the B cell surface to drive IgE isotype switching (*40, 41*). This mechanism does not explain the MBC2 phenotype as it is present in mice lacking IgE, and MBC2 cells lowly express CD21 (Fig. 2C). CD23 is also reported to restrain IgE production, a role that can be at least in part attributed to the involvement of CD23 in negatively regulating BCR signaling (*42, 43*). When considered together, the phenotype of MBC2s suggests polarization by IL-4, expression of markers known to inhibit IgE production, and a transcriptional program which retains these cells in the memory fate.

We found that MBC2s are the primary clonal relatives of the allergen specific IgE which emerges during the first month of allergen SLIT, providing strong human *in vivo* evidence that this population is a reservoir for memory IgE. Curiously, greater expression of IL-4Rα alone is not sufficient to explain the connection between MBC2s and memory IgE responses, as a recent study found that IgE CSR occurs at a similar rate between IL-4Rα^hi^ and IL-4Rα^lo^ naïve B cells (*44*). Baseline expression of εGLT in MBC2s could be interpreted as evidence of a predisposition to IgE isotype switching upon allergen encounter. Further, one hypothesis is that εGLT expression allows for spontaneous IgE isotype switching even in the absence of activation, which would result in tonic IgE BCR signaling that drives PC differentiation and serum IgE production (*45, 46*). Rare homeostatic MBC2 IgE isotype switching events may also explain the few cases (8/90,000 transcriptomes) of rearranged IgE in the MBC2 cluster, and the very low albeit significant levels of IgE VH transcripts found in most blood samples (present study and (*16*)). Such a pathway might explain the persistence of circulating IgE in allergic patients despite the short lifespan of IgE^+^ PCs. However, this hypothesis requires that the εGLT is expressed from the productively rearranged antibody locus, rather than the allelically excluded, non-productive antibody locus. The non-productive allele can support class switch recombination, which is preceded by GLT transcription (*47*). εGLT transcription from the non-productive allele most likely could not result in IgE isotype switching on the productive allele, and therefore may be better interpreted as a sign of type 2 polarization. It is not possible, in our dataset, to determine which of these two hypotheses is most accurate. Additionally, it is possible that the expression of antigen presentation and costimulatory machinery, and the downregulation of inhibitory receptors like CD32 are the features of MBC2s which drive their propensity to contribute to memory IgE responses. Additional research is needed to elucidate the features of MBC2s that predispose them to IgE isotype switching by comparison to other MBC subsets.

The identification of the MBC2 phenotype among allergen-specific cells provides a potential biomarker and target for therapy in allergic diseases largely mediated by IgE, such as allergic rhinitis and food allergy. Future research should examine the shifts in the MBC2 phenotype during allergen immunotherapy as protection begins, beyond the one-month time point examined here. IgG4 induction is associated with protection in allergen immunotherapy and is also highly represented among MBC2s (Fig. 1J). Elucidating the pathways which drive IgE versus IgG4 production from MBC2s is crucial to understand whether permanent reprogramming of allergen-specific memory is possible.

The interaction between humans and helminths has been a constant of human evolution. MBC2s may have emerged as rapid responders for protection against re-infection with helminths. In societies with lower parasite burden, these same cells are the ones which target allergens and drive allergic pathologies. MBC2s may similarly protect against venoms (*48*), and may drive other pathologies in which IgE has been implicated, such as autoimmunity and atherosclerosis (*49, 50*).

## Materials and Methods

### Patients

Birch allergic were recruited as part of clinical trial TT-04 (EudraCT 2015-004821-15) and sampled at the ALK Hoersholm site, Denmark (*51*). The trial was in accordance with the Declaration of Helsinki and in compliance with Good Clinical Practice guidelines as set by the International Conference on Harmonisation. Relevant national ethics committees and regulatory authorities approved the trial protocol and amendments.

HDM allergic and non-allergic donors were recruited as part of the ALK patient cohort at the ALK Hoersholm site, Denmark. Blood acquisition was approved by the Danish local Ethics Committee. 50 mL of blood was drawn from donors after which PMBCs were isolated using Lymphoprep density gradient medium (STEMCELL, 07861). Cells were washed three times and frozen in RPMI containing 30% FCS and 20% DMSO using a Planer Kryo 10 Series II controlled rate freezer (Planer). The cryotubes were subsequently moved to a −150 °C freezer.

Peanut allergic donors were recruited from the Allergy Clinic at the McMaster Children’s Hospital and non-allergic donors from the McMaster University and Hamilton community. Blood acquisition was approved by the Hamilton Integrated Research Ethics Board. Equal or less than 2.5% of total blood volume was drawn from donors. Human peripheral blood mononuclear cells (PBMCs) were isolated through density gradient centrifugation, filtered with 40 µm strainer (VWR-CA21008-949 (352340)) and resuspended in 10% dimethyl sulfoxide-supplemented FBS. Cyrogenic tubes were transferred to −80°C in Mr. Frosty Freezing Containers (5100–0001) before long-term storage in liquid nitrogen.

### Mice

Laboratory mice were housed at the Central Animal Facility at McMaster University in specific pathogen free conditions. All experimental procedures were approved by McMaster University’s Animal Research Ethics Board. C57Bl/6NCrl and Balb/cAnNCrl were purchased from Charles River. KN2 mice (Il4^tm1(CD2)Mmrs^; (*32*)) and MB1-Cre Bcl6^fl/fl^ (Cd79a^tm1(cre)Reth/EhobJ^ x Bcl6^tm1.1Mtto^; (*52*)) were provided by Dr. Irah King (McGill University) and bred in-house. IgE-deficient (*55*) mice were provided by Dr. Hans Oettgen (Harvard University). Mice were kept on a 12-hour light/dark cycle and given low-fat food and water *ad libidum*.

### Murine Models

Unless otherwise specified, mice were sensitized by a single intraperitoneal injection of 250 µg ovalbumin (Sigma; A5478) and 5 µg cholera toxin (List Labs; 100B) in PBS. Epicutaneous sensitization was performed by applying 200 µg of ovalbumin following shaving or tape-stripping the lower back (*13*). Intragastric sensitization was performed by four oral gavages of 1 mg of ovalbumin and 5 µg in 500 µl of PBS once a week for four weeks. Type 1 immunization was performed using 250 ug ovalbumin and 40 ug of CpG ODN 1826 (InvivoGen; vac-1826-1).

### Tetramer Construction and Enrichment

B cell tetramers were constructed as previously described (*56*). Monomeric antigens were biotinylated (ThermoFisher; A39257) at an optimized ratio which resulted in <1 biotin per molecule. Excess biotin was removed using an Amicon Ultra size exclusion column with an appropriate pore size for each molecule. Antigens were tetramerized by incubation at a >4:1 ratio of biotinylated protein with streptavidin-PE (Agilent; PJRS25) or streptavidin-APC (Agilent; PJ27S). Decoy tetramers were constructed by conjugating the same streptavidin-PE with Dylight594 (ThermoFisher; 46413), or streptavidin-APC with Dylight755 (ThermoFisher; 62279) and loading with an irrelevant protein; thereby ensuring all epitopes within the fluorescent backbone and biotin linker are excluded from the analysis. The following antigens were used: ovalbumin (Sigma; A5503), Ara h 2 (ALK Abelló), Bet v 1 (ALK Abelló) Spike receptor binding domain from native SARS-CoV-2 (Generous gift from Dr. Matthew Miller and Dr. Mark Larché, McMaster University).

Mouse antigen-tetramer enrichment was performed on pooled spleen and mesenteric lymph node cells from each mouse (100-200 million cells). Human antigen-tetramer enrichment was performed on purified PBMCs (80-100 million cells). Samples were stained with 1 µl of 1 µM decoy tetramer for 5 minutes at room temperature prior to adding 1 µl of 1 µM tetramer for 25 minutes on ice. After washing with 15 ml of FACS buffer, cells were incubated with 25 µl anti-PE (Miltenyi Biotec; 130-105-639) or anti-APC microbeads (Miltenyi Biotec; 130-090-855) for 15 minutes on ice. Bead-bound cells were positively selected through a magnet-mounted MACS LS column (Miltenyi Biotec; 130-042-401). In some experiments, PBMCs from multiple donors were pooled and then divided into two equal portions for enrichment with different antigens, allowing for a direct comparison.

### RNA Sequencing

#### 10x genomics

Switched memory B cells were isolated from human PBMCs from allergic and non-allergic donors by negative selection using the Switched Memory B Cell Isolation Kit, an LS Column, and a MidiMACS™ Separator (Miltenyi Biotec). Switched memory B cells were processed into single cell suspension for 10x single cell immune profiling (5’ gene expression + BCR). Cell suspensions (10.000 cells pr sample) were loaded according to 10x Genomics Manufacturer’s instructions for Single Cell A Chip to generate Gel Bead-In-Emulsion (GEMS, 10x genomics) using barcoded gel beads. RNA containing GEMS underwent barcoded cDNA synthesis. Subsequent library construction followed 10x Genomics 5’ Single Cell Immune Profiling Manufacturer’s Instructions. Libraries were constructed using the Chromium Next GEM Single Cell 5’ Library & Gel Beads Kit v1.0 (cat. No. 1000006) with additional Chromium Single Cell V(D)J Enrichment Kit, Human B Cell kit for BCR Libraries (cat. no. 1000016). Generated cDNA was quantified with Qubit 4 (Invitrogen) and quality assessment with Agilent 2100 Bioanalyzer (Agilent). Sequencing was then performed on Illumina NextSeq500.

#### SMARTSeq2

SMARTSeq2 library preparation was performed on single antigen-specific B cells single-cell sorted into PCR-grade RNase and DNase free 96-well plates. Plates were stored in −80°C prior to library preparation. Cells were lysed and reverse transcribed into cDNA. cDNA was then amplified and purified before quantification with Qubit 4 (Invitrogen) and quality assessment with TapeStation 4150 (Agilent). Library preparation for Illumina sequencing was completed with NovaSeq 6000 Reagent Kits (Illumina; 20028400). cDNA was fragmented, index labelled, amplified, and pooled prior to sequencing. Sequencing was then performed on Illumina NovaSeq 6000.

#### Bulk RNA Sequencing

Total RNA was isolated from the flowthrough of the negative selection of switched memory B cells in the six birch allergics at both baseline and at the 4 weeks of SLIT timepoint. The samples were sent to iRepertoire for heavy chain bulk sequencing.

### Flow Cytometry

Cryopreserved human PBMCs were thawed and enriched for antigen-specific cells as described. Cells were plated in 96 well U-bottom plates (up to 3 million/well). The cells were first resuspended in Human TruStain FcX (Biolegend; 422302) in 25 µl FACS buffer for 15 minutes on ice. 75 µl of extracellular staining cocktail was added for 30 minutes. Clones are dilutions are listed in Supplementary Table 1.

Mouse spleens were crushed through a 40 µm strainer using the plunger of a 3 ml syringe. Mouse mesenteric and inguinal lymph nodes were crushed between glass slides and passed through a 40 µm strainer. For antigen-enrichment experiments, splenocytes and mesenteric lymph node cells were pooled and processed for tetramer enrichment. Otherwise, red blood cells were removed from splenocyte preps using ACK lysis buffer (prepared in-house). Up to 3 million cells were plated into a 96 well U-bottom plate for staining. Cells were resuspended in FcBlock (Biolegend; 93; 101302) in 25 µl FACS buffer (2% FBS 1mM EDTA in PBS) for 15 minutes. 25 µl of extracellular staining cocktail was added for a final staining volume of 50 µl, and incubated for 30 minutes on ice. Antibody clones and dilutions are present in Supplementary Table 2; concentrations are recorded for the 50 µl staining volume. In experiments where anti-IgM and GL7 were used together, GL7 was stained in a separate extracellular staining step to avoid cross reactivity of the anti-mouse IgM antibody with the rat IgM GL-7 antibody.

### Cell Culture

25,000 ACK-lysed splenocytes from naïve C57Bl/6 mice were plated in 96-well U-bottom plates in 200 µl RPMI complete with 10% FBS, 1% penicillin/streptomycin, 1% L-Glutamine, 1x β-mercaptoethanol. Cultures were supplemented with 150 ng/ml anti-CD40 and 25 ng/ml IL-4 Cells were incubated in a 37°C 5% CO_2_ incubator. All steps were performed in sterile conditions. Flow cytometry was performed on day 5 of culture.

### Setup of HEK293 cells for expression of rIgEs

The cells used for expression of the rIgE clones construct were Freestyle 293-F cells (Thermofisher, cat no: K900001/510029). A 1 mL stock of 1*10^7 cells were rapidly thawed from −80°C in 37°C water bath and transferred to 19 mL prewarmed Freestyle™ 293 Expression Medium (Thermofisher, cat no: 12338018) and incubated at 37°C at 8% CO2 under orbital shaking at 125 rpm in a 125 mL Erlenmeyer flask. After 5 days the cells were spun down at 80g for 7 minutes at 20°C and media were changed to 30 mL prewarmed Freestyle™ 293 Expression Medium with 0,5% Penicillin-Streptomycin (Thermofisher, cat no: 15140122), vortexed vigorously for 20 seconds to break cell clumps, transferred to a new Erlenmeyer flask, and incubated at 37°C at 8% CO2 under orbital shaking. After additional five days the cells were spun down at 80g for 7 minutes at 20°C, split 1:3 in 30 mL pre-warmed Freestyle™ 293 Expression Medium with 0,5% Penicillin-Streptomycin, and then incubated at 37°C at 8% CO2 under orbital agitation. Four days after the cells were split 1:6 by previous stated centrifugal method, and five of these 1/6th splits were incubated in 40 mL prewarmed Freestyle™ 293 Expression Medium with 0,5% Penicillin-Streptomycin and incubated at 37°C at 8% CO2 under orbital agitation for 3 days before transfection.

### Biolayer interferometry kinetic profiles

For IgE all experiments were conducted on the Octet RED93e (Sartorius) at 30°C, 1000 RPM agitation, and 2 Hz frequency (high sensitivity setting). Data were processed with Data Analysis HT 11.1 software (Sartorius). High Precision Streptavidin (SAX) Biosensors were conditioned for a minimum of 10 minutes in kinetic buffer (DPBS, 0,1% BSA, 0,02% Tween-20, pH 7,4) before starting the experiment. Enzymatically biotinylated sdab026 (patent WO 2012’/175740 A1) with an AviTag™ (GLNDIFEAQKIEWHE) at the C terminal, were loaded onto the SAX biosensors at a concentration of 0,375 ug/mL for 180 seconds with a resulting signal of approximately 0,7 nm. Sensors were then dipped into kinetic buffer until a stable baseline were achieved. Then the sensors were dipped into hIgE mAbs diluted 1:10 in kinetic buffer for 300 seconds for an approximate signal of 2,5 to 3,5 nm. Sensors were then dipped into kinetic buffer until a stable baseline were achieved. Sensors were then dipped into the allergen of interest at a concentration 100 nM for 300 seconds of association followed by 300 seconds of dissociation in kinetic buffer. Regeneration down to immobilized sdab026 were achieved with 3×15 second pulses of 10 mm glycine (Sigma Aldrich) at pH 1,5, with a 15 second pulse in kinetic buffer in between each acid pulse. A stable baseline in kinetic buffer were achieved before the next set of hIgE mAbs were bound by immobilized sdab026.

### ELISA

A Nunc MaxiSorp™ flat-bottom 96 well plate (Thermofisher, cat no: 44240421) were coated overnight with 100 μL Birch pollen extract, Bet v 1, 10 ug/mL rabbit anti human IgE, or 2% BSA. Birch pollen extract was 50 μg/mL and Bet v 1 was 2,5 μg/mL. 2% BSA and 10 μg/mL rabbit anti human IgE were used as a negative and positive control, respectively. All the reagents were diluted in DPBS. After overnight incubation at 4°C with the coating reagents the wells were washed twice with ELISA wash buffer and blocked with 200 μL 2% BSA for 2 hours at room temperature and agitation. Following blocking the wells were washed three times with ELISA wash buffer and incubated for one hour at room temperature with 100 μL of the various rIgE’s supernatants diluted 1:10 in 2% BSA. After incubation with the rIgE’s the wells were washed four times with ELISA wash buffer and incubated with 100 μL of 1,3 μg/mL rabbit anti-human IgE-conjugated HRP (DAKO, cat no: P0295) for 1 hour at room temperature. The wells were washed four times in ELISA wash buffer, whereafter 100 μL TMB (Kem-En-Tec, cat no: 4380A) were added and the plates were shaken for 10 minutes exactly with no light exposure. After the 10 minutes the reaction were stopped with 100 μL 0,5 M sulphuric acid (VWR Chemicals, cat no: 30144.294). The plates were read at 450 nm on an EL808 Ultra Microplate Reader (BioTek Instruments)

### Data processing

#### 10x genomics

Cell ranger (v. 6.1.2) analysis was used to assemble the transcripts, alignment to human genomes and counting of aligned sequence reads for individual genes. The analysis was performed with human genome GRCh38-2020-A for gene expression (5GEX) and GRCh38-alts-ensembl-5.0.0 for BCR-vdj (VDJ). The 5GEX filtered_feature_bc_matrix from the output of cell ranger was imported into Seurat (version 4.1.1) for further analysis. We filtered out cells with less than 200 genes detected and genes that are only detected in less than 10 cells. We further filtered out cells with percentages of mitochondria genes more than 7%. Celldex (v. 1.4.0) and HumanPrimaryCellAtlasData (snapshotDate. 2021-10-19) was used to predict the cell phenotypes and used to remove non-MBCs. We manually removed two small impurities from megakaryocytes (PPBP+ PF4+ CAVIN2+) and from T cells (CD3E+ IL7R+ IL32+), that were not removed by the celldex filtering, leaving a total of 90,418 MBCs. The gene expression counts were normalized and scaled using the inbuild SCTransform function of Seurat. Integration features were selected using SelectIntegrationFeatures, however, IGHV, IGK, IGL genes were excluded from this list, since these dominated the subsequent clustering. The integrated object was analyzed by PCA (npcs = 50) and UMAP (dims 1:50), after which the cells were clustered based on Shared Nearest Neighbors using Euclidean distances (FindNeighbors, resolution 0.4). Differentially expressed genes were determined using the Wilcoxon Rank Sum test (FindAllMarkers). All upregulated differentially expressed genes greater than a threshold of 0.25 were used for gene ontology analysis using the Generic Gene Ontology Term Finder (https://go.princeton.edu/cgi-bin/GOTermFinder). The VDJ all_contig_annotations from the output of cell ranger resulted in 79,194 MBCs with identified isotypes. Of the 90,418 MBCs remaining after filtering of the 5GEX data, 73,258 (81%) were also identified in the VDJ pipeline.

#### SMARTSeq2

The sequencing results from the tetramer sorted MBCs were analyzed using the RNA-sequence aligner STAR (v. 2.7.10a). The resulting count matrices were loaded into Seurat, normalized and scaled, and mapped (FindTransferAnchors and MapQuery) onto the reference UMAP generated from the 90,418 MBCs from the 5GEX analysis.

B cell receptor sequences were reconstructed using a modified version of the BraCeR pipeline (docker pull nielsphk/bracer:1.3), originally created by Lindeman et al. 2018 (https://pubmed.ncbi.nlm.nih.gov/30065371/). Heavy and light chain sequences for a subset of identified cells, were used to generate plasmid constructs (Genscript) for recombinant expression of antibodies on a IgE background.

#### IgE coclustering

Bulk sequencing reads of six birch allergics were trimmed (first 3 bp of R1) with seqtk (v.1.3) and demultiplexed with ea-utils fastq-multx (1.02.772). Pair end reads were merged with PEAR (0.9.7) and the primer was trimmed with seqtk (19 bp at each end). Sequences were collapsed to unique sequences with fastx_collapser (fastx_toolkit-0.0.14) with counts. IgE annotated heavy chain sequences from bulk sequencing as well as consensus sequences from the VDJ analysis by 10x genomics were annotated with MiGMAP (v.0.9.8). Annotated sequences were combined per subject and clustered using changeo (dist=0.1, model=ham, v.0.3.1). Clusters containing both IgE reads from bulk (counts >10) as well as representatives from 10x VDJ were deemed to be coclustering. A subset of coclustering cells were selected for expression and recombinantly expressed in HEK293. All code is available at https://github.com/KoenigJFE/MBC2.

## Acknowledgements

We thank Gitte K. Koed and Jette Skovsgaard for excellent technical assistance and the University of Copenhagen Core Facility for Flow Cytometry and Single Cell Analysis for support. We thank McMaster University’s Centre for Advanced Light Microscopy, Dr. Joao Bronze de Firmino, and Dr. Jose Moran-Mirabal for access to analysis computers and support. We thank the McMaster Flow Cytometry Core, Hong Liang, and Minomi Subapanditha for access to analysis flow cytometers, experimental support, and flow sorting. We thank McMaster’s Central Animal Facility, Marion Corrick, Tim Ryan, and Dr. Eric Seidlitz for support in animal experimentation. We thank Dr. Justin J. Taylor (Fred Hutchinson Cancer Research Centre) for support and protocols for B cell tetramer construction, and Dr. Matthew Miller and Dr. Mark Larché (McMaster University) for providing RBD for tetramer construction. We thank Adam Wade-Vallance for input and feedback during manuscript preparation. We thank Claud Spadafora for support with visual design.

## Funding

Schroeder Foundation (MJ, SW, JFEK) Food Allergy Canada (MJ, SW, JFEK) ALK Abelló A/S (MJ, JFEK), Zych Family (MJ, SW) Satov Family (MJ, SW), Canadian Allergy Asthma and Immunology Foundation (MJ)

## Author contributions

Conceptualization: JFEK, NPHK, AP, KB, MJ, PSA

Formal Analysis: JFEK, NPHK, AP, KB, IH, DRG

Investigation: JFEK, NPHK, AP, KB, GL, DDL, AL, LHC, TW, AF

Visualization: JFEK, NPHK, AP, KB

Funding acquisition: JFEK, SW, MJ, PSA

Supervision: SW, MJ, PSA

Writing – original draft: JFEK, NPHK, KB

Writing – review & editing: JFEK, NPHK, AP, KB, IH, AL, LHC, DRG, MJ, PSA

## Competing interests

JFEK, SW, MJ receive funding from ALK Abelló A/S. PSA is on the advisory board of the Schroeder Allergy and Immunology Research Institute at McMaster University. AP, KB, IH, GL, DDL, AL, LHC, DRG, TW, AF declare that they have no competing interests.

## Data and materials availability

Code for this manuscript is available at https://github.com/KoenigJFE/MBC2. MBC atlas scRNA-seq transcriptomes, human Bet v 1-specific MBC transcriptomes, and mouse OVA-specific MBC transcriptomes will be deposited with a GEO accession prior to publication. All other data are available in the main text or supplement of the manuscript.

## Supplementary Text

### scRNA-seq reveals 21 human MBC clusters

The 90,418 sequenced MBCs formed 21 clusters that were detected in all of the participants included in the study (Fig. S1B-C). Clusters 3 and 12 are the two MBC2 clusters IGHE^-^ and IGHE^+^ clusters described in the main text.

Clusters 1, 11, and 13 are related clusters which closely align with the CD95^+^ memory cluster identified by Glass *et al* (*5*). These clusters express CRIP1 and CRIP2, CD99 and CD29 (ITGB1). They share HOPX expression with MBC2s, and some allergen-specific B cells and IgE-related cells in Figures 3 and 4 co-cluster in these populations. Indeed, the cluster most related to the MBC2 clusters in Euclidean space is cluster 11. Cluster 11 expresses IL-27B (EBI3), the serotonin receptor HTR3A, and the lymph node homing integrin CD62L (SELL). Cluster 13 is highly enriched in IgA2 MBCs, while cluster 11 and 13 equally express IgA and IgG isotypes.

Cluster 2 is the AREG^+^ cluster discussed in the main text. Cluster 20 is closely related to the AREG^+^ cluster. Both clusters are slightly enriched in IgA isotype MBCs and share expression of Amphiregulin and multiple repressor genes-CDKN1A, ZFP36, NR4A2, GADD45B (*57–59*). The major distinguishing feature between these two clusters is the very high expression of ID3 in cluster 20, a transcriptional regulator associated with repression of TCF4 and involved in germinal center responses. Clusters 7 and 9 are also related to cluster 2. Cluster 7 expresses greater Fos and Jun than cluster 2, while cluster 9 is differentiated by expression of the RNA helicase DDX5, mitochondrial genes CO1 and ND5 and ND6, and expression of the complement inhibitor CD55 (*60*).

Cluster 19 is the CD19^hi^ CD11c^hi^ cluster described by Glass et al and in the main body of the paper, a population also called atypical B cells. These cells have been reviewed elsewhere; in brief they show evidence of interferon signaling (IFI30) and type 1 polarization (ZEB2, TBX21). Cluster 19 cells are most similar to CD95^+^ memory cells, consistent with Glass *et al*. While this population is transcriptionally distant from most other populations, some cells in clusters 2 and 4 show transcriptional similarity to cluster 19, resulting in their relatively lower Euclidean distance.

Cluster 4 is highly enriched in IgG2 MBCs, and differentially expresses Galectin-1 (LGALS1), TCF4, and FCLR2 (*61*). This population also shares transcriptional similarity to cluster 5, a population defined primarily by expression of J chain, which is partially involved in IgM and IgA multimerization, and is involved in binding to pIgR for mucosal IgA and IgM transport (*62*). J chain expression is associated with plasma cells, but cluster 5 cells do not express plasma cell associated transcription factors like Prdm1 and XBP1 (data not shown).

Clusters 0, 8, and 16 are related clusters whose differential expression is largely based in the downregulation of many genes expressed by other clusters. Cluster 0 expresses suppressive genes like GAS5, TPT1, and TXNIP, as well as the antisense non-coding RNA FAM30A which is located in the IgH locus (53, *54*). Cluster 8 is defined by higher JUN and WDR74 expression. Cluster 16 differentially expresses genes associated with interferon signaling, STAT1, IRF1, and IRF9. Cluster 18 is a distinct population that highly expresses multiple interferon stimulated genes (ISGs, STAT1, STAT6, IFIs, MX1, LY6E).

Cluster 10 is a population of MBCs which have a transcriptional signature associated with B cell receptor signaling (NR4A1, EGR2), activation (CD69, CD83) and lymph node homing (CCR7). Cluster 7 has similar gene expression to cluster 10, expressing fewer genes of B cell receptor signaling, but greater FOS.

Clusters 14 and 17 both express genes mapping to the IGHM and IGHD locus. For cluster 17, which we reported as the IgM^+^ IgD^+^ cluster in the main text, this is confirmed by the VDJ amplification in Supplemental Fig. 2D. However, cluster 14 VDJ reads are primarily non-IgM isotypes, suggesting that these cells may be expressing IgM germline transcripts. Cluster 17 expresses naïve B cell markers TCL1A, YBX3. The other major defining feature of cluster 14 was the absence of the antisense non-coding RNA FAM30A (*54*). Cluster 15 is most similar to cluster 14 in Euclidean space, its primary distinguishing factor is the expression of STAG3, a stromal antigen with a known role in meiosis.

Cluster 6 cells differentially express multiple ribosomal proteins, and some mitochondrial DNA. This population is expressed in all individuals but may represent cells undergoing apoptosis.

---

**Fig. S1:**
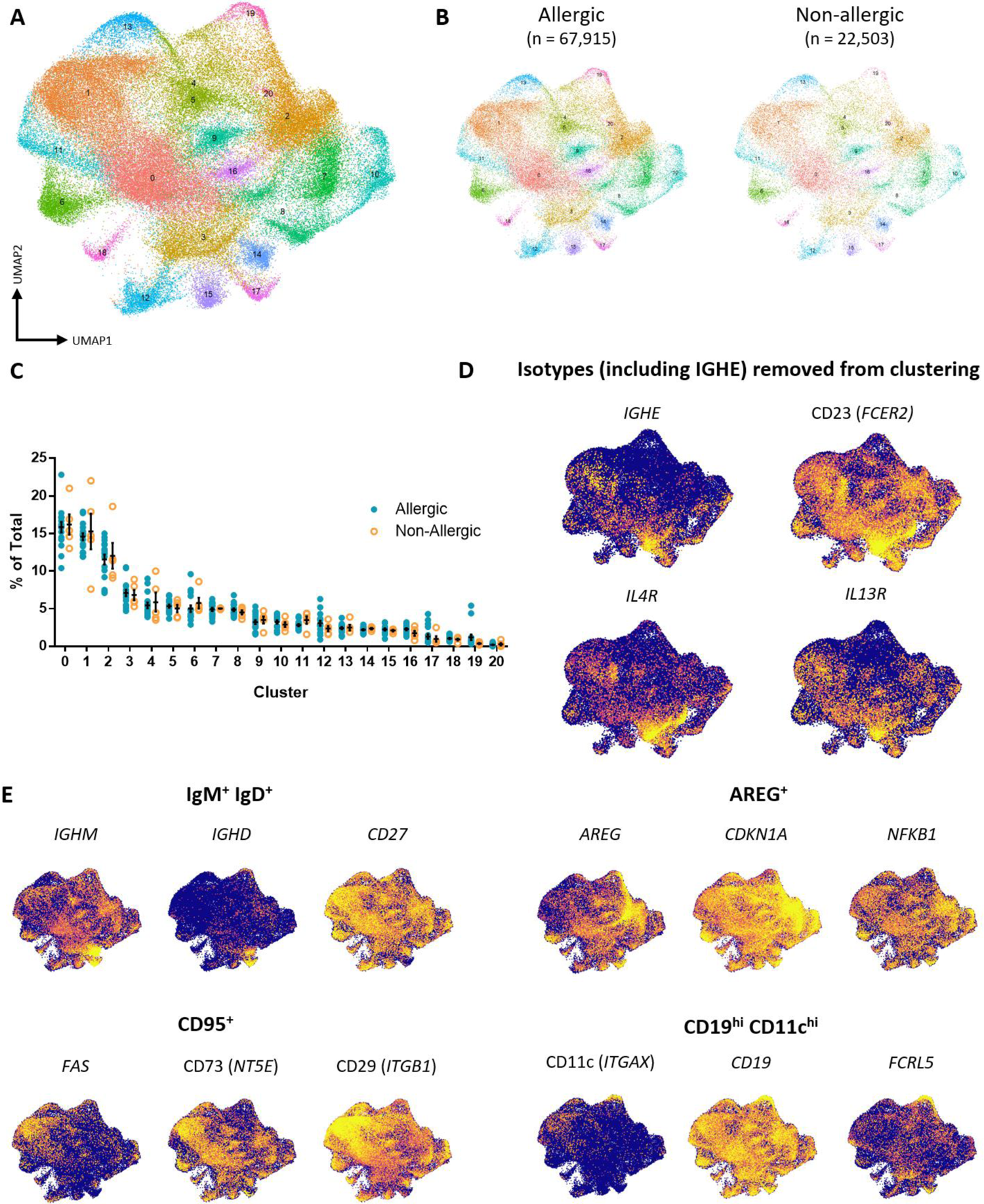
scRNA-seq MBC atlas revealed 21 clusters that were detected at similar levels in all participants. (**A**) UMAP plot of transcriptomes from human MBCs from peripheral blood. (**B**) UMAP plot separated by allergic status of participants. (**C**) Frequency of each cluster from total MBCs. (**D**) UMAP plot when MBCs isotype is not considered in dimensionality reduction, colorized by expression of MBC2-associated genes. (**E**) Expression of defining features of the four clusters selected for comparison against MBC2s.

**Fig. S2:**
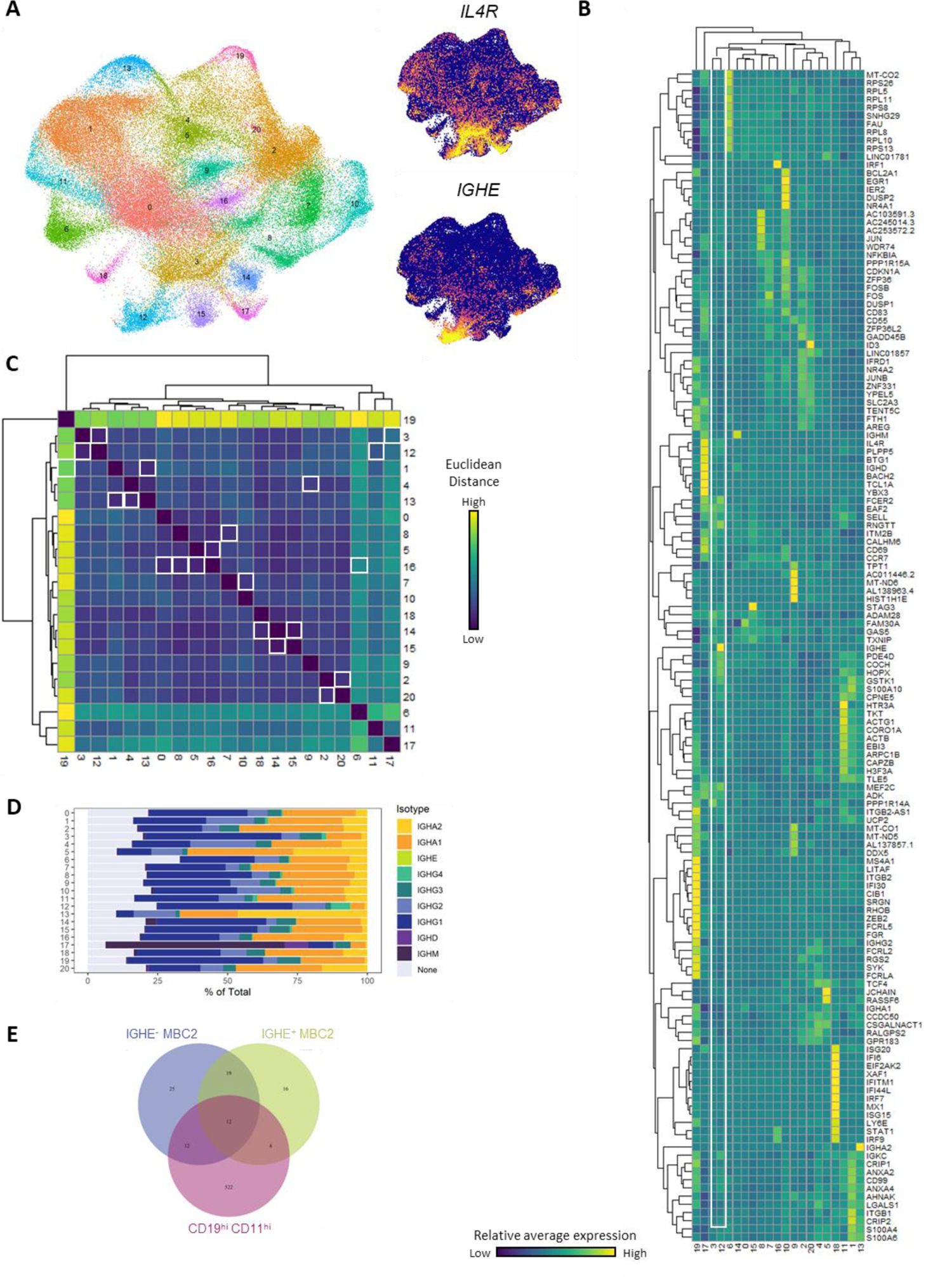
Analyses from Figure 1 including all clusters. (**A**) UMAP plot of human MBC clusters, *IL4R*, and *IGHE* expression for reference. (**B**) Heatmap of top 10 differentially upregulated genes for each cluster. (**C**) Euclidean distance between clusters. Lowest distance for each cluster is denoted with a white outline. (**D**) Isotype identified from amplified VDJ sequences by cluster. (**E**) Overlap of differentially upregulated genes between the IGHE^-^, IGHE^+^, and CD19^hi^ CD11c^hi^ clusters.

**Fig. S3:**
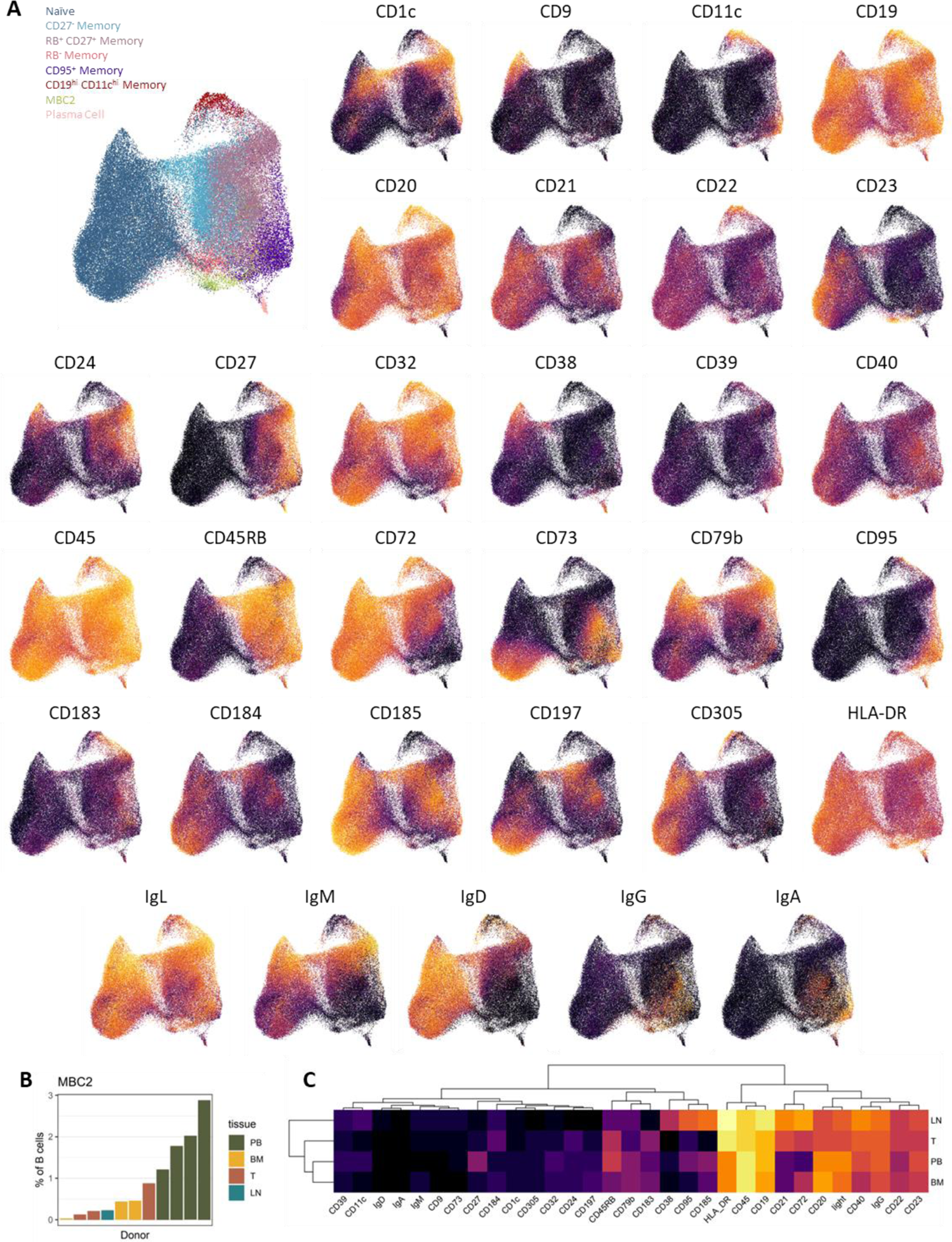
Comprehensive surface phenotypic profiling of human MBC2s. (**A**) UMAP plots colorized by marker for mass cytometry data on human MBCs from Glass *et al.* (**B**) Frequency of MBC2s among B cells from the peripheral blood (PB), bone marrow (BM), tonsil (T), and lymph nodes (LN). (**C**) Heat map of surface expression differences between MBC2s across tissues.

**Fig. S4:**
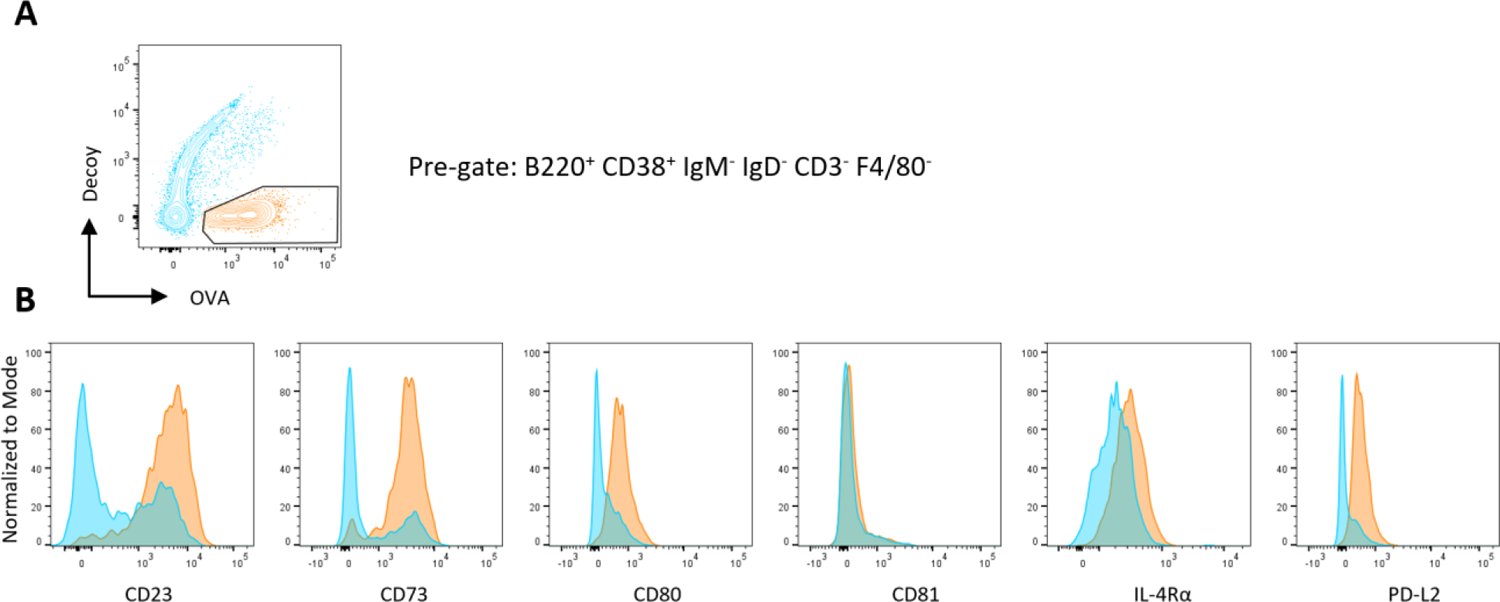
Surface phenotype of mouse MBC2s. WT mice were immunized with OVA and CT i.p. and pooled OVA-specific spleen and mesenteric lymph node cells were analyzed at 1-month post-sensitization. (**A**) Representative gating of OVA-specific and non-allergen-specific cells. (**B**) Histograms of markers of MBC maturity on OVA-specific (orange) and non-allergen-specific (blue) MBCs.

**Fig. S5:**
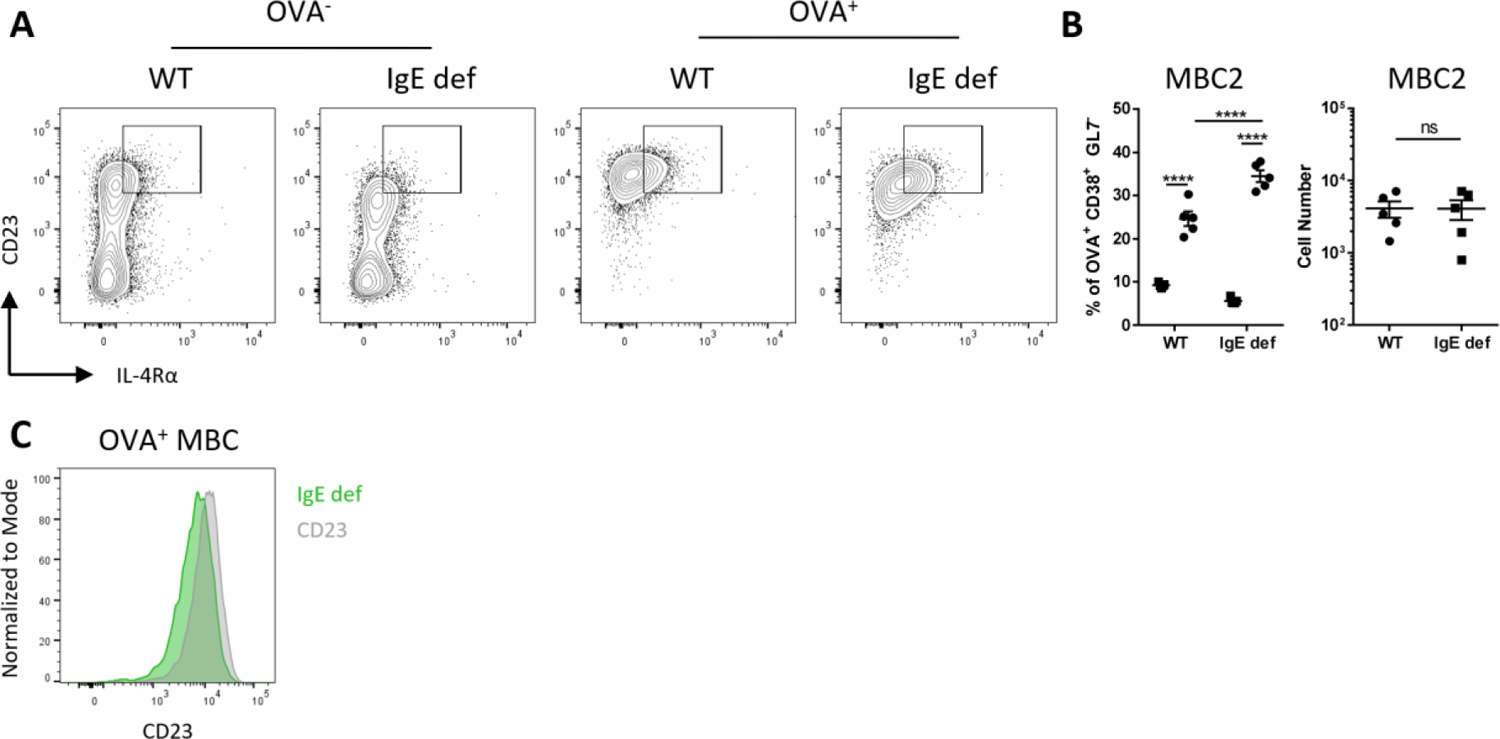
MBC2 differentiation does not require IgE. IgE deficient (IgE def) and WT mice were sensitized i.p. with OVA and CT. (**A-B**) Concatenated flow plots, frequency and cell number of switched OVA-specific memory B cells from pooled spleen and mesenteric lymph nodes. (**C**) MFI of CD23 expression on OVA-specific MBCs in IgE def and WT mice. * p < 0.05 ** p < 0.01 *** p < 0.001 **** p < 0.0001 via Mann-Whitney test.

**Table S1. (separate Excel file)**

Differentially expressed genes for all 21 clusters identified in the scRNA-seq MBC atlas.

**Table S2.** Human Flow Cytometry

| **Supplementary Table 2: Human Flow Cytometry** |  |  |  |  |
| --- | --- | --- | --- | --- |
| **Marker** | **Fluorophore** | **Concentration** | **Clone** | **Source (Catalogue Number)** |
| CD14 | BV711 | 1:100 | M5E2 | Biolegend (301838) |
| CD16 | BV711 | 1:100 | 3G8 | Biolegend (302044) |
| CD19 | PerCP-Cy5.5 | 1:20 | SJ25C1 | Biolegend (363016) |
| CD23 | BV421 | 1:20 | EBVCS-5 | Biolegend (338522) |
| CD27 | BV605 | 1:20 | O323 | Biolegend (302830) |
| CD3 | BV711 | 1:100 | OKT3 | Biolegend (317328) |
| CD38 | PECy7 | 1:100 | HB-7 | Biolegend (356608) |
| IgD | AF700 | 1:20 | IA6-2 | Biolegend (348230) |
| IgM | BV650 | 1:20 | MHM-88 | Biolegend (314526) |
| IL-4Rα | PE | 1:20 | GO77F6 | Biolegend (355004) |
| Viability Dye | Aqua | 1:1000 |  | Thermofisher (L34957) |

Antibodies used for human flow cytometry.

**Table S3.**
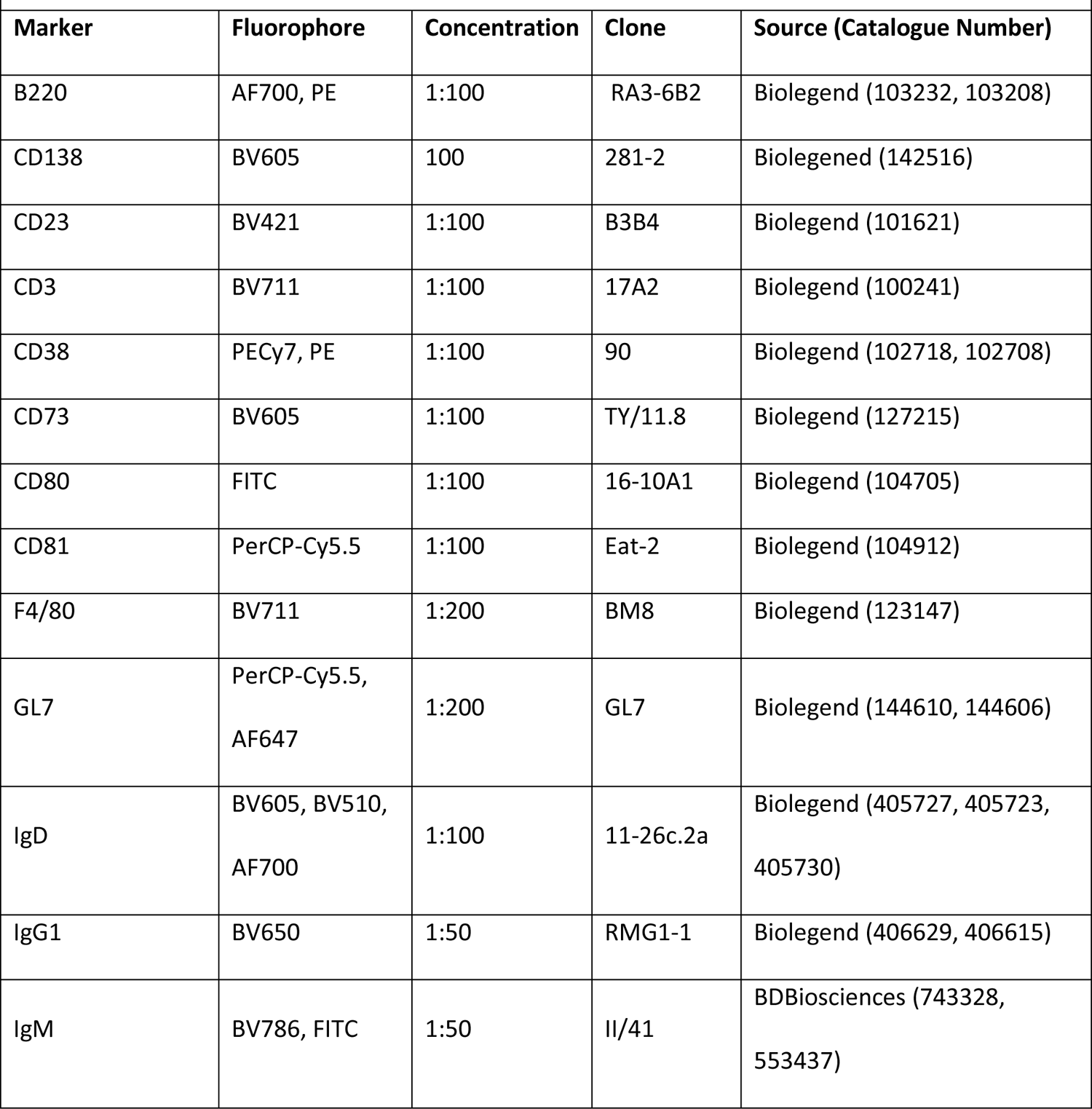
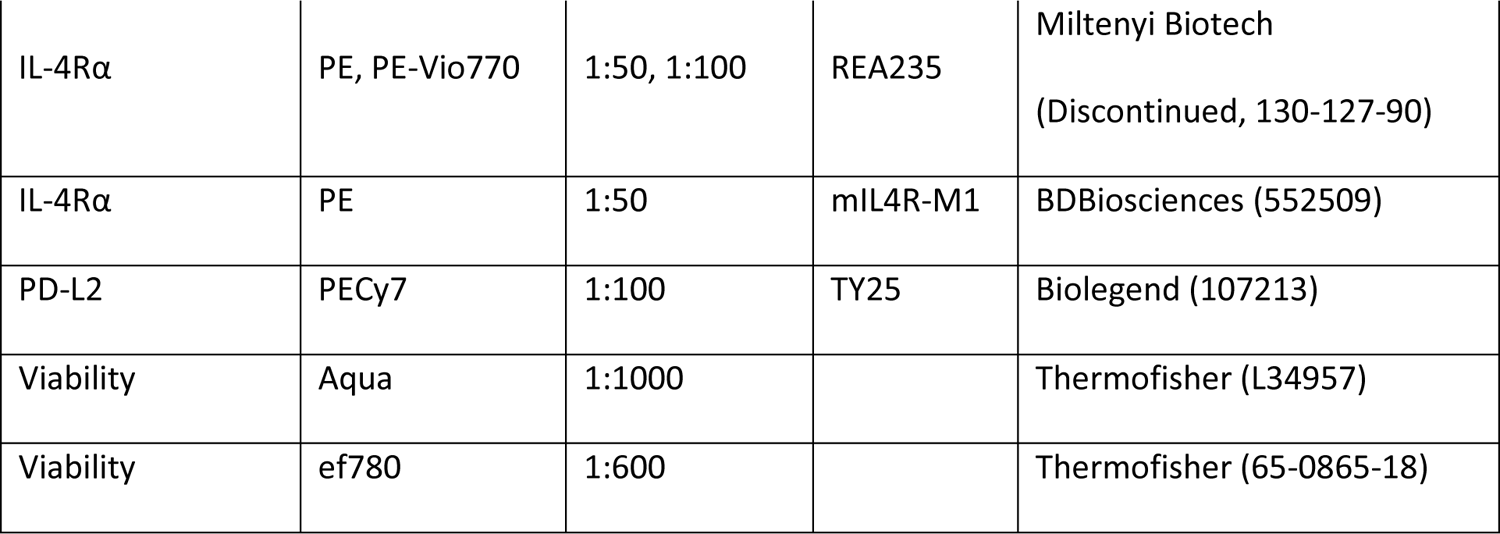
Antibodies used for mouse flow cytometry. Supplementary Table 3: Murine Flow Cytometry

| **Supplementary Table 3: Murine Flow Cytometry** |  |  |  |  |
| --- | --- | --- | --- | --- |
| **Marker** | **Fluorophore** | **Concentration** | **Clone** | **Source (Catalogue Number)** |
| B220 | AF700, PE | 1:100 | RA3-6B2 | Biolegend (103232, 103208) |
| CD138 | BV605 | 100 | 281-2 | Biolegend (142516) |
| CD23 | BV421 | 1:100 | B3B4 | Biolegend (101621) |
| CD3 | BV711 | 1:100 | 17A2 | Biolegend (100241) |
| CD38 | PECy7, PE | 1:100 | 90 | Biolegend (102718, 102708) |
| CD73 | BV605 | 1:100 | TY/11.8 | Biolegend (127215) |
| CD80 | FITC | 1:100 | 16-10A1 | Biolegend (104705) |
| CD81 | PerCP-Cy5.5 | 1:100 | Eat-2 | Biolegend (104912) |
| F4/80 | BV711 | 1:200 | BM8 | Biolegend (123147) |
| GL7 | PerCP-Cy5.5,<br>AF647 | 1:200 | GL7 | Biolegend (144610, 144606) |
| IgD | BV605, BV510,<br>AF700 | 1:100 | 11-26c.2a | Biolegend (405727, 405723,<br>405730) |
| IgG1 | BV650 | 1:50 | RMG1-1 | Biolegend (406629, 406615) |
| IgM | BV786, FITC | 1:50 | II/41 | BDBiosciences (743328,<br>553437) |
| IL-4Rα | PE, PE-Vio770 | 1:50, 1:100 | REA235 | Miltenyi Biotech<br>(Discontinued, 130-127-90) |
| IL-4Rα | PE | 1:50 | mIL4R-M1 | BDBiosciences (552509) |
| PD-L2 | PECy7 | 1:100 | TY25 | Biolegend (107213) |
| Viability | Aqua | 1:1000 |  | Thermofisher (L34957) |
| Viability | ef780 | 1:600 |  | Thermofisher (65-0865-18) |

## Notes

### Competing Interest Statement

JFEK, SW, MJ receive funding from ALK Abello A/S. PSA is on the advisory board of the Schroeder Allergy and Immunology Research Institute at McMaster University. AP, KB, IH, GL, DDL, AL, LHC, DRG, TW, AF declare that they have no competing interests.

